# Emergent epistasis mediates the role of negative frequency-dependent selection in bacterial strain structure

**DOI:** 10.1101/2025.05.02.651819

**Authors:** Martin Guillemet, Sonja Lehtinen

**Affiliations:** Department of Environmental System Science, Institute for Integrative Biology, ETH Zürich, Zürich, Switzerland; Swiss Institute of Bioinformatics, Lausanne, Switzerland, Department of Computational Biology, University of Lausanne, Lausanne, Switzerland

## Abstract

Strain structure is a well-documented phenomenon in many pathogenic and commensal bacterial species, where distinct strains persist over time exhibiting stable associations between genetic or phenotypic traits. This structure is surprising, particularly in highly recombinogenic species like *Streptococcus pneumoniae*, because recombination typically breaks down linkage disequilibrium, the non-random association of alleles at different loci. Allelic diversity is a necessary condition of strain structure. Recent work suggests that multi-locus negative frequency-dependent selection (NFDS) acts to maintain diversity across bacterial genomes. Here, using modelling and genomic analysis, we show that multi-locus NFDS also shapes bacterial strain structure through emergent epistatic effects. We develop models of two NFDS mechanisms – metabolic niche differentiation and competition-colonisation trade-offs – and show that epistasis emerges readily in these models. Notably, both models generate frequency-dependent epistasis. Unlike classical epistasis, this acts to either reinforce or abolish existing strain structure, making observed allele associations contingent on the evolutionary history of the population. We then use a dataset of over 3000 *S. pneumoniae* genomes to test our model predictions, and make observations consistent with emerging epistatic effects on gene associations. Our results extend and generalise previous work on the role of antigen-specific acquired immunity (a diversity-maintaining mechanism) on strain structure. Overall, this works contributes to a better understanding of the evolutionary processes shaping the structure of bacterial populations, which is central to predictive modelling of multi-strain pathogens

## Introduction

Many pathogenic and commensal bacterial species exist as distinct strains that are stable through time, a phenomenon referred to as strain structure. These strains can vary significantly in their phenotypic traits, such as immunological characteristics, antibiotic resistance, competitive abilities, or average duration of carriage [1, 2, 3, 4, 5]. For instance, certain strains of *Streptococcus pneumoniae* are more likely to cause invasive disease, impacting vaccine design and implementation [3]. Similarly, hyper-invasive clonal complexes in *Neisseria meningitidis* have been observed to persist and spread globally over several decades [1, 2]. Strain structure is characterised by strong patterns of linkage disequilibrium (LD) over the whole genome, meaning that pairs of alleles tend to be found together more or less often than expected [6]. This leads to stable associations between phenotypic traits. For example, resistances to different antibiotics tend to co-occur [5], and pneumococcal strains characterised by a longer duration of carriage are more likely to be resistant [4]. Characterising the mechanisms leading to stable associations between phenotypes of interest is important for understanding the ecology of these traits and the dynamics of bacterial strains.

In the absence of recombination, strain structure is a fundamental consequence of clonal reproduction. However, recombination breaks down strain structure by shuffling alleles between different genetic backgrounds, thus eroding non-random associations between loci. Over time, in absence of other factors, recombination is expected to lead to linkage equilibrium. The recombination rate required for strain structure to be eroded in the presence of genetic drift is not entirely clear. Nevertheless, especially in highly recombinogenic species such as *S. pneumoniae* [7], we would expect stable strain structure to require active maintenance by selective forces – more specifically epistasis.

Epistasis refers to the interaction between genes at different loci, where the effect of one allele (or gene) on a phenotype is affected by the presence of one or more other alleles: the combined effect is different from the sum of the individual allele effects. Epistasis can stem from molecular interactions, for example with compensatory mutations that alleviate the cost of antibiotic resistance genes [8]. It can also arise from effects on distinct life-history traits because of how these life-history traits combine in the expression of fitness. For instance, antibiotic resistance is expected to be more advantageous to strains that have a long duration of carriage [4, 9]. More broadly, however, the extent to which observed allele associations across bacterial genomes arise from epistasis and the mechanisms giving rise to this epistasis are not fully understood.

Here, we explore the role of multi-locus negative frequency-dependent selection (NFDS) in giving rise to epistasis and in shaping bacterial strain structure. The work is motivated by the detection of wide-spread NFDS acting across bacterial genomes [10, 11, 12]. NFDS is a form of balancing selection which promotes allelic diversity through rare alleles having a selective advantage and thus maintains accessory genes at intermediate frequencies. The mechanisms giving rise to this wide-spread NFDS are not fully characterised, but plausible candidates include antigen-specific acquired immunity [13, 14, 15], metabolic niche differentiation [16] and trade-offs related to direct bacterial competition [17]. Note that here we use the term NFDS consistently with previous literature [10], which, in the case of acquired immunity may not be strictly accurate [18] because the fitness of variants depends on their historical rather than current frequency. Previous modelling work has suggested NFDS acts to maintain strain structure when combined with asymmetric recombination favouring gene deletion over gene acquisition, through the emergence of outbreeding depression [19]. In this work, we suggest NFDS also shapes strain structure more directly, through emergent epistasis.

Previous models of multi-locus NFDS [10, 19] have generally assumed selection acting on multiple loci combines additively. However, it is not clear this assumption holds, particularly for functionally related loci. For example, NFDS on antigenic alleles arises because antigen-specific acquired immunity makes transmission to previously colonised hosts more difficult, disadvantaging historically frequent antigens [20]. When considering multiple loci, how this NFDS combines across the loci depends on the relative strength of the immune response to strains carrying one versus two previously encountered antigens. There is no *a priori* reason to assume this should be additive. Indeed, there is considerable theoretical work exploring the effects of different assumptions on the associations between antigenic alleles [21, 22, 23, 24, 25, 26].

Here, we expand and generalise these ideas by showing how multi-locus NFDS can – depending on how it combines across loci – either abolish or maintain strain structure through emerging frequency-dependent effects on allele combinations. We first explore two epidemiological models with different mechanisms of NFDS to demonstrate how these effects arise. We then develop a phenomenological model that captures and abstracts the key insights. Finally, we test specific predictions from this model in a dataset of more than 3000 pneumococcal isolates, showing patterns consistent with the frequency-dependent effects we predict.

## Results

### A Simple Model of Multi-locus Negative Frequency-Dependent Selection

We begin by revisiting linkage disequilibrium in the multi-locus NFDS model from Corander et al. [10], in which NFDS combines additively across loci. Our goal here is not to provide new results, but rather to build intuitions and introduce concepts and terminology important to understand the rest of the paper.

This model considers a population of bacteria with two bi-allelic loci, resulting in four possible genotypes *G* = {*ab, Ab, aB, AB*}. The dynamics of each genotype are governed by logistic growth with additive effects from NFDS at each locus. The growth rate (in the absence of recombination) of the genotype with alleles *i* and *j* are given by:

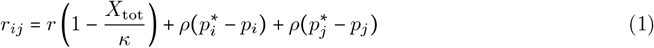

Here *r* is the intrinsic growth rate, *κ* is the carrying capacity, *p*_*x*_ is the frequency of allele 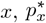 is the allele frequency favoured by selection and *ρ* is the strength of NFDS. Under this model, NFDS acts independently on each allele, favouring alleles that are below their optimum frequency and disadvantaging those above it. This mechanism ensures that allele frequencies are maintained at an intermediate frequency, but is not expected to maintain strain structure in the presence of recombination [19].

To formalise this idea, we quantify LD using *D*:

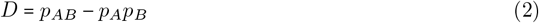

Thus, *D* = 0 indicates random association between alleles, while positive *D* indicates over-representation of *AB* and *ab*, and negative *D* indicates over-representation of *Ab* and *aB*. Note that we use the metric *D* for derivations, but prefer to use the metric *D*^′^ in figures as it is more readily interpretable and always bounded by − 1 ≤ *D*^′^≤ 1 independently of allele frequencies, see Methods. To understand the dynamics of LD, it is helpful to also define the individual selection coefficient on allele A (*w*_*A*_) and an epistatic term (*w*_*AB*_) as:

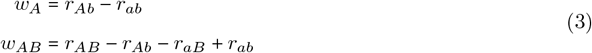

When individual allele frequencies are at equilibrium (*w*_*A*_ = *w*_*B*_ = 0) and introducing recombination, the dynamics of LD are given by [26]:

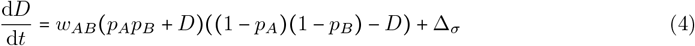

where Δ_*σ*_ is the net effect of recombination in the population on *D*. In the case of unbiased recombination, this is of sign opposite to current LD [27]. Due to the bounds on the values of D, the term multiplying *w*_*AB*_ is ≥ 0, so the sign of *w*_*AB*_ drives the build-up of negative or positive LD.

Turning back to our model of multi-locus NFDS, when allele frequencies are optimal, the growth rate is the same for all genotypes (Equation 1), and thus epistasis *w*_*AB*_ = 0. In other words, in absence of recombination, this model predicts neither increasing nor decreasing effects on LD. Introducing recombination into the model (see Methods), we get Δ_*σ*_ = − *σD*. Recombination drives *D* towards 0 over time, confirming the model does not sustain strain structure under this recombination scheme.

### Metabolic niche Model

We now turn to epidemiological models of multi-locus NFDS in order to understand what assumptions about NFDS combining across loci mean biologically, whether this is necessarily additive, and the effect these assumptions have on strain structure. We begin by considering a model in which NFDS arises from within-host competition being stronger when strains are similar. Here we frame this as arising from metabolic niche overlap, although other effects such as strain-specific immunity could also give rise to the same model structure.

#### Model description

Again, we consider a two-locus, two-allele model (*G* = { *ab, Ab, aB, AB* }). The full model structure is given in the Methods and the one locus case is represented in Figure 1 (see Supplementary Figure S1 for the two-locus case). Briefly, hosts can be uncolonised (*S*), colonised with a single strain (*I*_•_) or co-colonised with two strains (*I*_•,•_). Colonisation is cleared at rate *γ*, hosts become colonised through density-dependent transmission at baseline rate *β*_0_. Onward transmission of each strain from a co-colonised host is scaled down by a factor *q* compared to singly colonised hosts, reflecting reduced density within the host. A resident strain inhibits co-colonisation by incoming strains, modifying the transmission rate to already colonised hosts by an efficiency parameter *k*:

**Figure 1:**
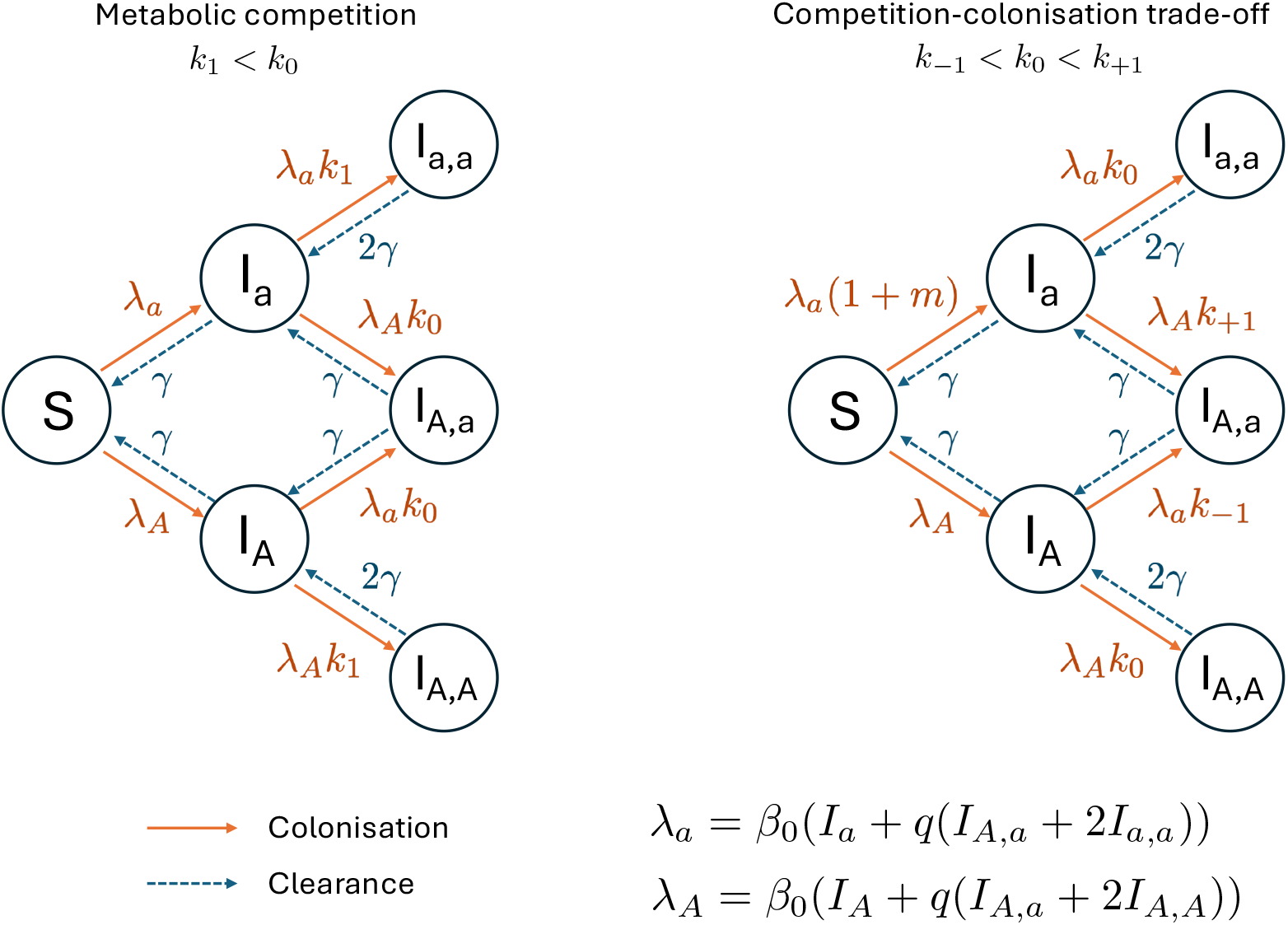
NFDS models of metabolic niche or competition-colonisation trade-offs. We represent the two epidemiological model for the case of one bi-allelic locus. In the metabolic niche model (left), co-colonisation rates are reduced when the resident and incoming strains share the same allele, modeled by parameters *k*_1_ < *k*_0_. In the competition-colonisation model (right), allele a is “colonising”, leading to an increased rate of primary colonisation *m*, while the competitive allele A leads to higher rates of co-colonisation on average. Co-colonisation rates are dependent on the difference in number of competitive alleles between the incoming and the resident strain Δ through the parameters *k*_Δ_. *λ*_*a*_ and *λ*_*A*_ represent respectively the partial forces of infection of allele *a* and *A*, taking into account that strains in co-colonisation have reduced transmission efficiency through parameter *q*. Solid orange lines represent colonisation and dashed blue lines represent clearance.

- *k*_0_ If the incoming strain differs from the resident strain at both loci (i.e. no shared alleles)
- *k*_1_ If the incoming strain shares one allele with the resident strain
- *k*_2_ If the incoming strain shares both allele with the resident strain

These parameters represent metabolic niches wherein different alleles provide access to different resources, with *k*_0_ > *k*_1_ > *k*_2_. Strains sharing metabolic pathways (i.e. with the same allele at a locus) have a high niche overlap, competing for the same resources within the host, while strains with different metabolic capabilities (i.e. with different allele at a locus) can coexist within a host more readily. This gives rise to NFDS at each locus, driving allele frequencies to 0.5 at each locus (see Supplementary Note 1 for an analysis of the NFDS in the one-locus case).

#### Metabolic niche differentiation leads to frequency-dependent effects on genotypes and strain structure

To understand the dynamics of LD in this model, we derive the expression for epistasis *w*_*AB*_ (Equations (3)). Assuming *q* = 1 for simplicity:

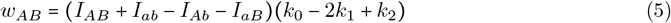

This expression indicates that epistasis is dependent on both the strain composition of the population and parameters relating to metabolic competition. The strain density term can be decomposed into the sum of the density of strains associated with positive LD (*AB, ab*), minus the sum of the density of strains associated with negative LD (*Ab, aB*). Thus, in this model, epistasis acts to reinforce or abolish existing LD, depending on the sign of the competition term.

The competition term (*k*_0_ − 2*k*_1_ + *k*_2_) reflects how NFDS combines across loci:

**If k**_**1**_**= (k**_**0**_ + **k**_**2**_**)/2**, NFDS combines additively across loci. Compared to co-colonising a host carrying an identical strain (i.e. two shared alleles, efficiency *k*_2_), each additional divergent allele provides the same additional benefit. In this case, epistasis is zero and there is not effect on strain structure.

**If k**_**1**_< **(k**_**0**_ + **k**_**2**_**)/2**, NFDS combines super-additively (Figure 2a, dashed line). Compared to co-colonising a host carrying an identical strain (*k*_2_), one divergent allele (*k*1) provides a small benefit relative to two divergent alleles (*k*_0_). The biological interpretation here is that both loci are important: one private resource (*k*_1_) facilitates co-colonisation only to a limited extent compared to both resources being private (*k*_0_). The (*k*_0_ − 2*k*_1_ + *k*_2_) term is positive: epistasis will therefore act to reinforce existing LD. This produces bi-stability (Figure 2b and Figure S3a): LD reaches either *D*^′^= ± 1 at equilibrium without recombination, depending on the initial sign of *D*.

**Figure 2:**
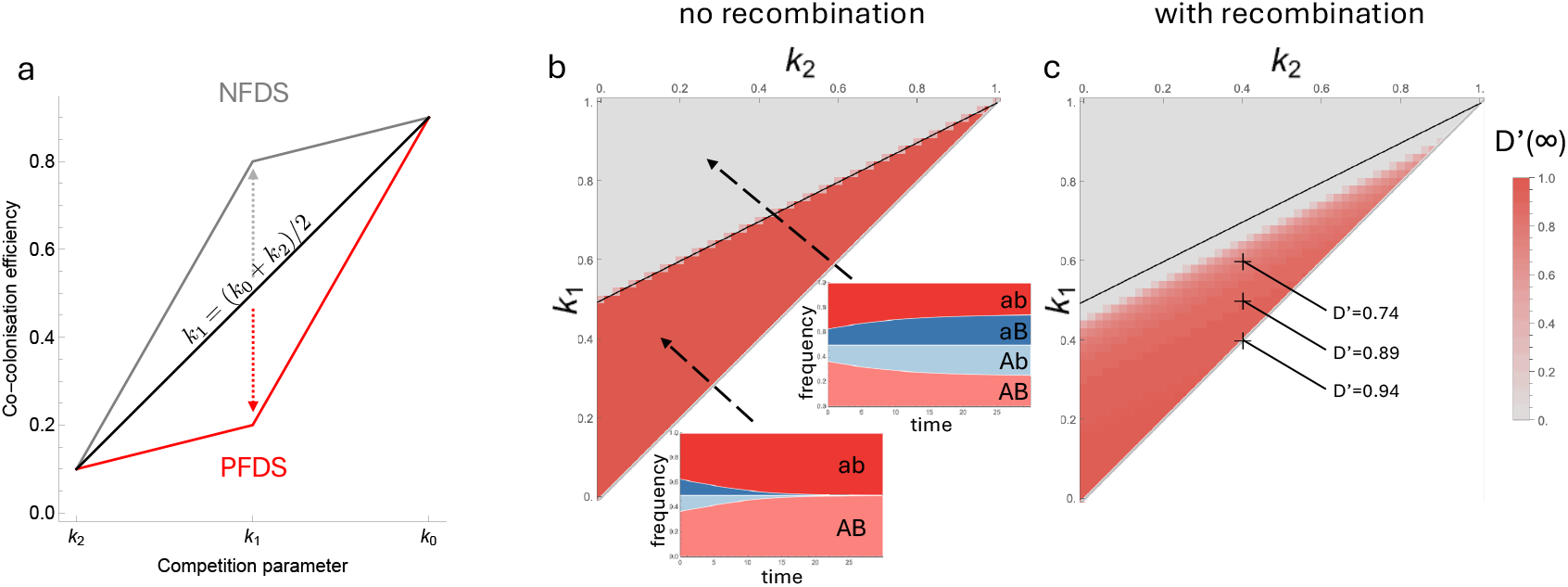
Different competition regimes lead to maintenance or abolishment of strain structure. In (a), we show how different geometries for the parameters competition efficiencies *k*_0_, *k*_1_ and *k*_2_ influence epistasis and therefore strain structure according to eq. (5). Equilibrium *D*′ (t=5000) is shown as a function of the competition parameters *k*_1_ and *k*_2_ in the absence (b) and the presence (c) of recombination. Müller plots in panel (b) show the dynamics of the cumulative frequencies of all genotypes through time, depending on the competition parameters, starting for an initial LD of *D* = 0.1. Some specific values of *D*^′^ are noted on panel *c*. Parameter values used: *β*0 = 2, *b* = 4, *γ* = 2, *d* = 1, *k*_0_ = 1, *q* = 0.5, (b,c) *k*_1_ = 0.6, *k*_2_ = 0.4 and in (c) *σ* = 0.02

**If k**_**1**_> **(k**_**0**_ + **k**_**2**_**)/2**, NFDS combines sub-additively (Figure 2a, solid line). Compared to co-colonising a host carrying an identical strain (*k*_2_), one divergent allele (*k*1) provides a large benefit relative to two divergent alleles (*k*_0_). The biological interpretation here is that there is redundancy between the loci: one private resource (*k*_1_) facilitates co-colonisation to a large extent, and the benefit of the other additional resource is limited (*k*_0_). The (*k*_0_ − 2*k*_1_ + *k*_2_) term is negative: epistasis will therefore act to abolish existing LD, driving *D*^′^ to zero at equilibrium (Figure 2b and Figure S3a).

In other words, NFDS combining super-additively across loci gives rise to positive frequency dependent selection (PFDS) on pairs of non-overlapping genotypes (i.e. *AB* and *ab*, or *Ab* and *aB*). This reinforces existing strain structure. On the other hand, NFDS combining sub-additively across loci gives rise to NFDS on genotypes. This abolishes strain structure, even in the absence of recombination. The model is only neutral with respect to LD when NFDS combines precisely additively. These results are robust to relaxing the assumption *q* = 1 as long as *q* does not become too small, a regime where co-colonised hosts are considerably less infectious overall than singly colonised host (*q* = 0.1 for the parameter set we use, see Figure S4).

Introducing recombination into the model (see Methods) accelerates the decay of LD when genotypes are under NFDS (Figure S5). When genotypes are under PFDS, the addition of recombination has a two-fold effect (Figure 2c). Firstly, the parameter space in which strain structure is maintained decreases: in the presence of recombination, strain structure is abolished unless PFDS is strong enough to overcome its effects. Secondly, in the parameter space in which strain structure is maintained, *D*^′^< 1 at equilibrium due to the balance between the effects of epistasis and recombination.

In summary, the metabolic niche model demonstrates that epistatic interactions resulting from NFDS processes combining across loci can lead either to abolishment or maintenance of strain structure, depending on the geometry of the generated fitness effects.

### Competition-Colonisation Trade-off Model

We now explore a second epidemiological model, in which NFDS arises through a different mechanism: a competition-colonisation trade-off. This type of NFDS arises when one allele provides an advantage in transmitting to uncolonised hosts (‘colonisation’) while the other leads to better rates of co-colonisation to already colonised hosts (‘competition’). This gives rise to NFDS because more colonisers decreases the ratio of uncolonised hosts to single infected hosts thus favouring competitors, and *vice versa*. This type of trade-off can arise from bacteriocin systems [28] or metabolic genes adapted to high vs low-nutrient (i.e. already colonised) environment.

#### Model Description

We use the same co-colonisation SIS framework, which is described in the methods, and the one locus case is represented in Figure 1 (see Supplementary Figure S1 for the two-locus case). In this model, each locus has two alleles: a ‘colonising’ allele (*a* and *b*), which increases transmission to uncolonised hosts (*S*), and a ‘competitive’ allele (*A* and *B*), which increases transmission to already colonised hosts. We assume additive effects of the colonising alleles such that the primary colonisation rates of the *i* genotype, *β*(*i, S*) is:

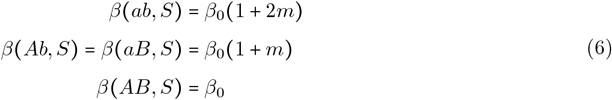

where (*β*_0_) is the baseline transmission rate and *m* is benefit associated with the colonising allele. Transmission to already colonised hosts is modulated by *k*_Δ_, with Δ the difference in the number of competitive alleles between the incoming (*i*) and the resident (*j*) strains:

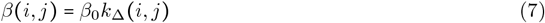

where Δ ∈ {−2, −1, 0, 1, 2}. For example *k*_Δ_ (*Ab, AB*) = *k*_−1_ and *k*_Δ_ (*AB, ab*) = *k+*_2_, with 0 ≤ *k*_−2_ < *k*_−1_ < *k*_0_ < *k*_+1_ < *k+*_2_ ≤ 1. Note that unlike the metabolic niche model, the alleles are not symmetric. However, genotypes *Ab* and *aB* are phenotypically equivalent. We explore this model and the resulting NFDS in case of a single bi-allelic locus in Supplementary Note 2 .

#### Competition-colonisation can maintain a variety of strain structure patterns

The epistatic term *w*_*AB*_ for *q* = 1 is given by (see Methods):

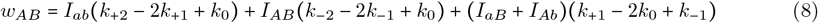

As before, this expression can be decomposed into *k*-parameter terms multiplying the densities of strains. For each *I*_*i*_ term, the corresponding *k*-parameter term captures information about how effectively genotypes can co-colonise host already carrying strain *i*. More specifically, each term reflects the impact of additional competitive alleles on the co-colonisation efficiency. For example, the first term (*k*_+2_ − 2*k*_+1_ + *k*_0_) represents the difference between i) the incoming strain having two vs one competitive allele (*k*_+2_ − *k*_+1_) and ii) the incoming strain having one vs no competitive alleles (*k*_+1_ − *k*_0_). Thus, as before, these terms capture how effects combine across loci. Again, this can be represented by the geometry of the parameter curve (Figure 3a): negative terms correspond to the sub-additive effects (the second competitive allele has a smaller effect than the first) and a concave curve; positive terms correspond to super-additive effects (the second competitive allele has a greater effect than the first) and a convex curve.

**Figure 3:**
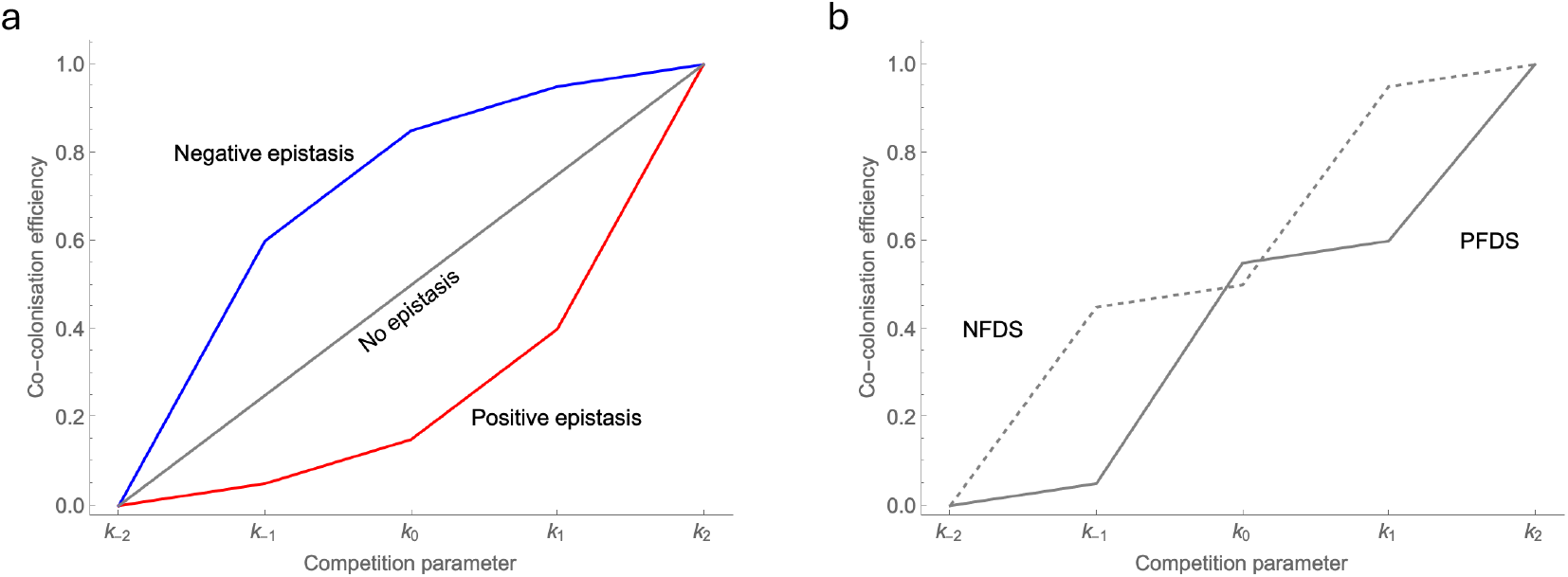
The geometry of competitive interactions shapes epistasis and strain structure in a competition-colonisation trade-off model. (a) A linear relationship between all the competition parameters *k*_Δ_ leads to additivity in how fitness effects combine across loci and thus no epistasis, see eq. (8). However a concave or a convex geometry will lead to epistasis of constant sign. (b) Step like geometries produce epistasis which sign is frequency-dependent. The solid line scenario leads to PFDS wherein the most frequent pair of non-overlapping strains (*D*+ or *D* − strains) is favoured. In contrast, the dashed line geometry leads to NFDS at the genotype level, favouring the least frequent pair of strains and thus abolishing strain structure. See Figure S7 and S8 for equilibrium LD from simulations.

The consequences of how these effects combine across loci are more complex than in the metabolic niche model. If all terms are positive (i.e. the entire curve is convex), *w*_*AB*_ is also always positive: competitive alleles are always most beneficial when on the same genome, leading to positive LD (i.e. *ab* and *AB* strains). Conversely, if all terms are negative (i.e. the entire curve is concave), *w*_*AB*_ is also always negative: competitive alleles are always most beneficial when not on the same genome, leading to negative LD (i.e. *Ab* and *aB* strains).

The model gives rise to frequency-dependent epistasis when the *k*-parameter terms are not all of the same sign (Figure 3b): in this case, the overall effect on *w*_*AB*_ depends on strain frequencies. For example, if the terms multiplying *I*_*ab*_ and *I*_*AB*_ (i.e. strains associated with positive LD) are positive and the term multiplying the *I*_*aB*_ and *I*_*Ab*_ (i.e. strains associated with negative LD) is negative, *w*_*AB*_ will have the same sign as the current LD. In other words, the model will give rise to PFDS on genotypes, re-enforcing existing LD. Conversely, if the terms multiplying *I*_*ab*_ and *I*_*AB*_ are negative and the term multiplying *I*_*aB*_ and *I*_*Ab*_ positive, *w*_*AB*_ will have the opposite sign to current LD, giving rise the NFDS on strain structure and driving LD to zero.

To explore a diversity of possible geometries, we illustrate in Figure S7 and S8 the equilibrium LD for a variety of parameter sets, as described in Supplementary Note 3. These figures show the robustness of the results for *q* = 1 and *q* = 0.5. Note however that we adjust the value of the colonising benefit *m* to maintain an equilibrium frequency of 0.5. This is because a low value of *q* disproportionately affects the competitive alleles, which are more likely to be found in co-colonising strains. Overall, depending on how the effects of individual alleles combine across loci, competition-colonisation tradeoffs can generate i) constant epistasis generating and maintaining structure, ii) PFDS on pairs of non-overlapping genotypes reinforcing pre-existing structure or iii) NFDS at the genotype level, abolishing structure.

### A Simple Model of Multi-locus NFDS with Epistasis

In light of these insights, we revisit the initial model of multi-locus NFDS and extend it to show that a simple model can reproduce the strain structure patterns observed in complex mechanistic models. We modify the growth rate expression to add deviation from additivity for the combining of NFDS across loci. First, as seen in the competition-colonisation trade-off model, pairs of alleles may be intrinsically beneficial or costly when together in the same genome, leading to constant sign epistasis *ϵ*_*ij*_. Secondly, both the metabolic and competition-colonisation model gave rise to frequency dependent effects on LD. We represent this effect by introducing the additional term −*ρ*_2_*p*_*ij*_. This term is subtracted so that the interpretation of *ρ*_2_ is the same as *ρ*: negative frequency-dependence when *ρ*_2_ > 0 and positive frequency-dependence when *ρ*_2_ < 0. This gives the following growth rate for genotype *ij*:

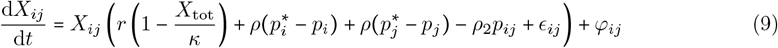

With this modification, we can reproduce a range of behaviours (Figure 4) observed in the mechanistic models. For *D*^′^ values, we get the range of possible values from −1 to 1 when constant epistasis is mixed with NFDS on genotypes (*ρ*_2_ > 0) with a single stable equilibrium. We also recover bi-stability when there is PFDS on genotypes (*ρ*_2_ < 0), which can overwhelm the effect of constant epistasis depending on the relative value of constant epistasis *ϵ*. We recover the same effects of recombination on equilibrium LD (see Supplementary Figure S9) as we found in mechanistic models: it reduces absolute LD for all parameter sets, and also shifts the threshold delimiting the parameter sets leading to a single stable equilibrium or to bi-stability (Figure S9).

**Figure 4:**
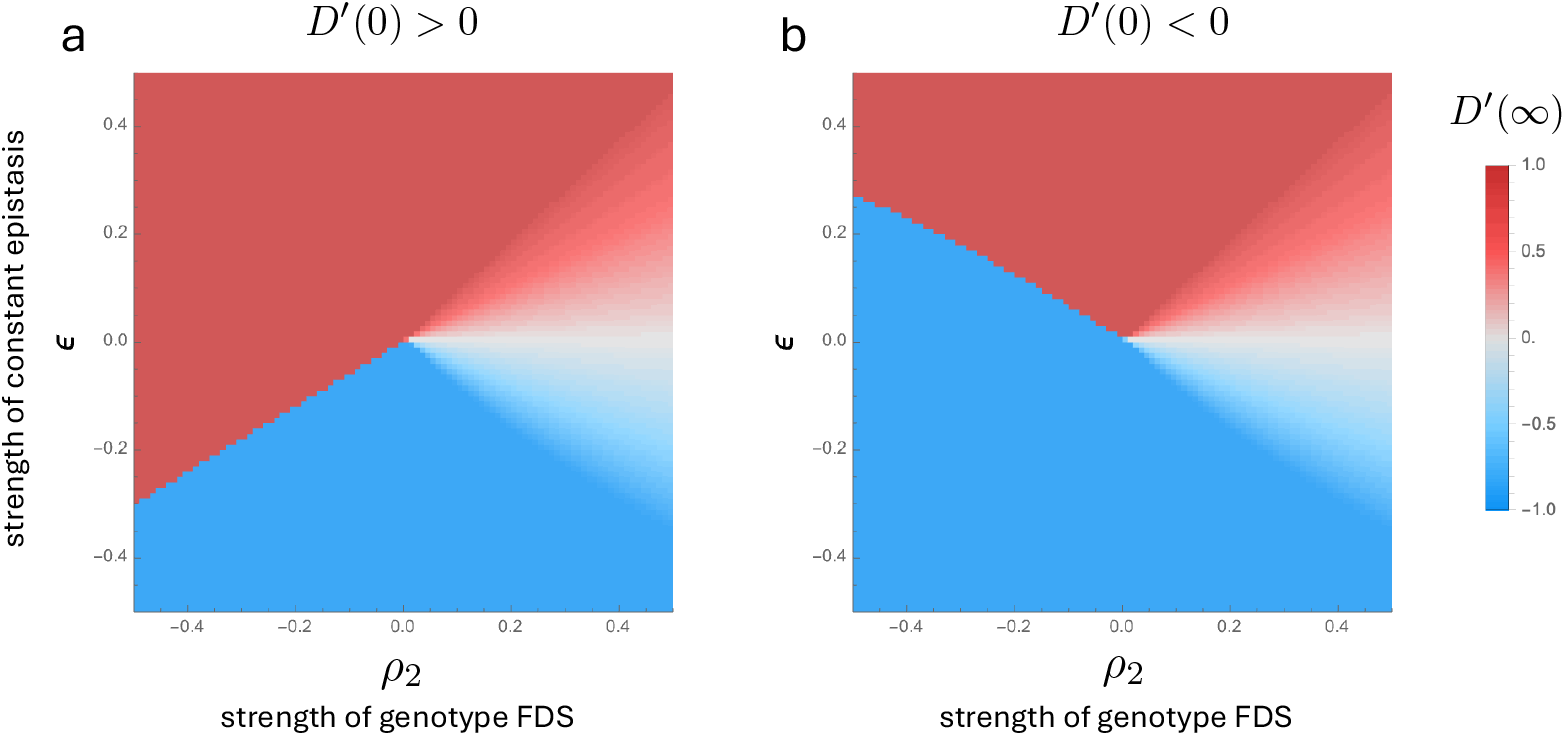
A simple multi-locus NFDS with constant sign and frequency-dependent epistasis reproduces varied strain-structuring effects. Equilibrium *D*′ from eq. (9) is shown as a function of the constant epistasis parameter *ϵ* and genotype frequency-dependent epistasis parameter *ρ*_2_ starting from a (a) positive or a (b) negative linkage disequilibrium *D*′(0), in the absence of recombination. See Figure S9 for the case with recombination. Parameter values used: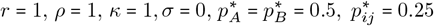.

This simple model demonstrates that the essential features of the more complex mechanistic models can be captured by incorporating frequency-dependent and constant epistatic effects into a multi-locus NFDS framework.

### Evidence for Frequency-dependent Effects on Strain Structure in *S. pneumoniae*

They key prediction from our models is that NFDS acting across multiple loci can give rise to frequency-dependent effects on strain structure. This prediction is difficult to test because, depending on parameter values, these effects can act to either abolish or reinforce strain structure. Furthermore, LD may also arise through other constant epistatic interactions between genes, or stochastic effects such as drift and hitch-hiking. Simply observing LD is thus not enough to discriminate between the mechanism suggested by our modelling and other possible causes. However, constant epistasis and PFDS make different predictions about LD *across* populations. Under constant epistasis, the same association is always favoured and the sign of LD must thus be the same across populations. PFDS, however, acts to reinforce existing structure; the sign of LD at equilibrium therefore depends on initial conditions. Pairs of genes with strong LD of different signs across different populations would therefore be consistent with PFDS, but not other forms of epistasis.

To test this idea, we examine patterns of linkage between 900 intermediate frequency genes across over 3000 pneumococcal genomes from three different locations: Thailand, Massachusetts and Southampton [29, 30, 31]. To alleviate confounding by stochastic events, we exclude genes which have been gained or lost fewer than 30 times on the whole *S. pneumoniae* tree. The dataset is represented in Figure S10.

We begin by analysing the functional categorisation of pairs of genes in LD (defined as | *D*^′^| > 0.6). We find that pairs of genes sharing a function are more likely to be in LD than random gene pairs (t-test: t=2.834, df=11.359, P-value=0.0158, Figure 5a). Assuming epistasis, whether frequency-dependent or constant, is more likely to arise between functionally related genes, this suggest patterns of linkage in this dataset are at least partly driven by epistasis rather than demographic events. Genes involved in energy production and conversion are particularly likely to show strong LD with other genes in the same category. Genes in the translation category (which notably includes integrases and transposases) show the strongest overall linkage, potentially reflecting the role of mobile genetic elements in LD.

**Figure 5:**
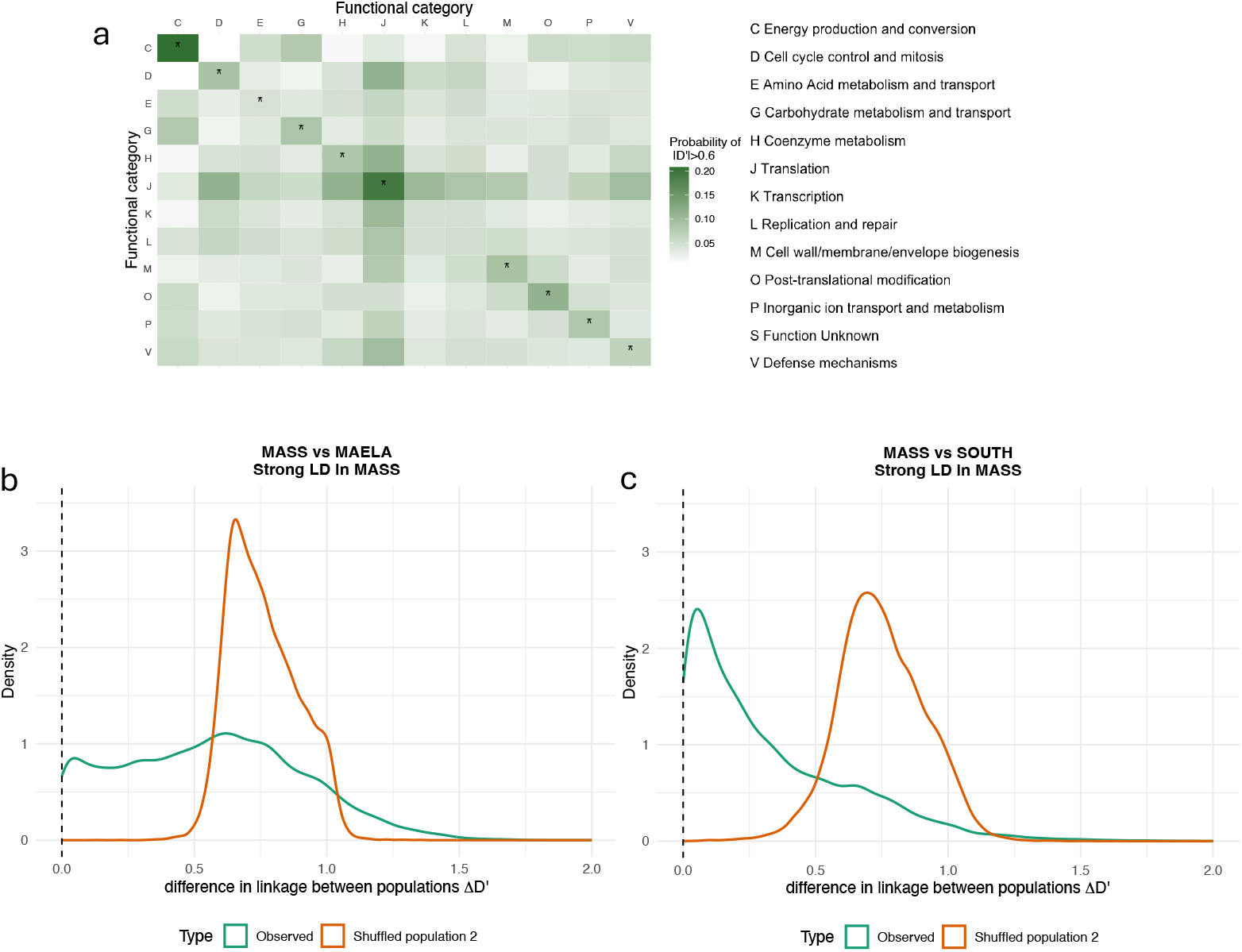
Observed patterns of linkage disequilibrium in several populations. (a) shows in the whole dataset, according to the KEGG annotation of a focal gene, the probability to be linked (| *D*^′^| > 0.6) with any given gene. An asterisk on the diagonal signifies that there is significantly more linkage for that combination (two genes with a shared KEGG category) compared to the rest of the row/column (one gene of that category with genes of different categories), removing the linkage with category J which is driven by mobile genetic elements. In (b,c),we study the distributions of absolute difference of LD across populations. We show for gene pairs highly linked in the Massachusetts dataset ( | *D*^′^| > 0.6) the distribution of change in linkage compared to (b) the MaeLa dataset or (c) the Southampton dataset. The green line shows the distribution of observed changes, and the orange line shows the distribution when the gene presence/absence data is permuted in the second population. See Supplementary Figure S12 for all comparisons and S11 for the representation of *D*^′^ of individual pairs across populations.

We then examine LD patterns across multiple populations, assessing the extent to which LD is conserved across populations. More specifically, for the set of gene pairs showing strong linkage in one population ( |*D*^′^| > 0.6), we look at the LD for each pair in the other populations, and compute the distribution of absolute change in LD between the focal and each of the other two populations (Figure 5b and 5c). We find that on average, the strong linkage is conserved across populations (defined as |*D*^′^| > 0.6 and same sign LD) for 30.6% of genes pairs, while it is reversed (defined as |*D*^′^| > 0.6 but opposite sign LD) for 2.7% of gene pairs. Linkage conservation is more frequent for more closely related populations like Southampton and Massachusetts (44.2%), whereas more isolated populations like Maela and Southampton show more frequent reversal (4.5%). To test the statistical significance of these results, for each comparison between populations, we permute the second population (keeping individual gene frequencies constant but shuffling the genomes each gene appears in) and recalculate the proportion of LD conservation and reversal (see Methods). Repeating this permutation 1000 times generates a null distribution under a model in which average LD is zero in the second population. Across all populations pairs, both LD reversal and conservation are significantly more frequent than expected based on the permuted populations (one-tailed test: p-value < 0.001 in all cases, Tables 1 and 2). Observations of large scale LD conservation are consistent with both constant and frequency-dependant epistasis. However, LD reversal is incompatible with constant epistasis and potentially indicates PFDS and contingency on the distinct evolutionary history of each population.

**Table 1:** Conservation of LD across populations and time.

| Comparison | Focal Population | Strong LD Pairs | Observed Conservation Events | Mean expected Conservation Events | *p*-value |  |
| --- | --- | --- | --- | --- | --- | --- |
|  |  |  |  |  | One-tailed | Two-tailed |
| maela-south | maela | 25 962 | 4230 | 0.008 | 0.001 | 0.001 |
|  | south | 11 087 | 4230 | 13.069 | 0.001 | 0.001 |
| mass-maela | mass | 9827 | 3190 | 6.190 | 0.001 | 0.001 |
|  | maela | 19 263 | 3190 | 0.006 | 0.001 | 0.001 |
| mass-south | mass | 20 595 | 8555 | 7.395 | 0.001 | 0.001 |
|  | south | 18 922 | 8555 | 11.684 | 0.001 | 0.001 |
| pre-post vaccine | 2001 | 25 745 | 11 650 | 551.469 | 0.001 | 0.001 |
|  | 2007 | 24 887 | 11 650 | 132.268 | 0.001 | 0.001 |

The table shows linkage analysis results between population pairs. Columns represent: compared populations, which focal population was analysed (Focal Population), number of strong linkage pairs analysed (Strong Pairs), observed conservations, mean permuted conservations (Permuted Mean), and p-values for one-tailed and two-tailed tests. Permuted means represent average conservations in null distribution. All p-values < 0.001 are shown as 0.001, and indicate that the observed differences are outside of the distribution of 1000 permutations.

**Table 2:** Reversal of LD across populations and time.

| Comparison | Focal Population | Strong LD Pairs | Observed Reversal Events | Mean expected Reversal Events | *p*-value |  |
| --- | --- | --- | --- | --- | --- | --- |
|  |  |  |  |  | One-tailed | Two-tailed |
| maela-south | maela | 25 962 | 447 | 0.005 | 0.001 | 0.001 |
|  | south | 11 087 | 447 | 4.383 | 0.001 | 0.001 |
| mass-maela | mass | 9827 | 172 | 2.625 | 0.001 | 0.001 |
|  | maela | 19 263 | 172 | 0.008 | 0.001 | 0.001 |
| mass-south | mass | 20 595 | 81 | 2.592 | 0.001 | 0.001 |
|  | south | 18 922 | 81 | 1.996 | 0.001 | 0.001 |
| pre-post vaccine | 2001 | 25 745 | 15 | 108.050 | 1.000 | 0.001 |
|  | 2007 | 24 887 | 15 | 23.082 | 0.976 | 0.084 |

The table shows linkage analysis results between population pairs. Columns represent: compared populations, which focal population was analysed (Focal Population), number of strong linkage pairs analysed (Strong Pairs), observed reversals, mean permuted reversals (Permuted Mean), and p-values for one-tailed and two-tailed tests. Permuted means represent average conservations in null distribution. All p-values < 0.001 are shown as 0.001, and indicate that the observed differences are outside of the distribution of 1000 permutations.

In principle, LD maintained through PFDS on strain structure might not be robust to strong perturbations. We investigate this by examining changes to LD patterns following the rollout of the pneumococcal conjugate vaccine in Massachusetts between 2001 and 2007 (Figure 5d), which targetted a subset of serotypes. In pre- and post-vaccination populations, consistent linkage was retained in 46 % of pairs, but the number of linkage reversal events was not higher than expected in the permutation-based null model. In fact, with 2001 as the focal population, the number of observed reversal events was significantly lower than expected (respectively 15 and 108 out of 25745 pairs, two-tailed test, p-value<0.001).

## Discussion

In this work, we show that NFDS acting across bacterial genomes [10, 11, 12] is expected to shape bacterial strain structure. Phenomenological models of multi-locus NFDS typically assume that the effects of NFDS combine additively across different loci [10, 19]. Here, we develop two mechanistic models of NFDS – within-host niches and competition-colonisation trade-off – and show that there is no *a priori* reason to assume effects combine additively. Deviating from this assumption leads to emergent epistatic effects. Under some parameter regimes, the competition-colonisation model gives rise to constant-sign epistatic effects, which predict stable associations between loci. More interestingly, both models also give rise to emergent frequency-dependent effects on strain structure. Negative frequency-dependence actively abolishes strain structure, while positive frequency-dependence acts to reinforce existing strain structure. The direction of associations maintained by positive frequency-dependent epistasis can therefore be in either direction, and is determined by historical contingency.

We examine patterns of association between intermediate frequency genes in more than 3000 pneumococcal genomes, from three different geographical populations. We find that strong positive or negative LD is more common than expected between functionally related genes, suggesting these associations are driven by epistatic rather than stochastic or demographic effects. Furthermore, we find that genes in strong positive or negative LD in one population are more likely than expected to be in strong positive *or* negative LD in another population. While conserved associations between populations is consistent with both constant and positive frequency-dependence epistasis, reversal is only consistent with positive frequency-dependence. Taken together, the analysis is therefore consistent with emergent positive frequency-dependent epistasis playing a role in maintaining strain structure. An important caveat here is that reversed associations were much rarer than conserved ones. Conserved associations are not informative in assessing the extent to which positive-frequency dependent epistasis shapes strain structure. It is also worth noting that these results require us to assume a null model and the choice of appropriate null model is not trivial. We use a permutation-based approach which assumes an average LD of 0. While this is standard when assessing the significance of LD, more realistic null models, explicitly incorporating estimated recombination rates and clonality, would allow more nuanced hypothesis testing.

While both our theoretical and empirical results suggest the frequency-dependent effects we find play a role in shaping strain structure, assessing the extent to which these effects play a role in observed patterns of LD is not trivial. Firstly, predicted frequency-dependent effects can act to either reinforce or maintain strain structure depending on how effects combine across loci. Empirical estimates of variation in colonisation efficiency are feasible (e.g. [32]); estimating how these effects are influenced by multiple loci requires large, sequenced surveys of bacterial colonisation.

Secondly, NFDS is hypothesised to maintain intermediate frequencies for genes across a wide range of functions, including immunity, metabolism, direct competition and antibiotic resistance [10]. We hypothesise that the emergence of frequency-dependent epistasis from multi-locus NFDS is a general effect. This work focuses on two mechanisms: within-host niches, primarily conceptualised as arising from metabolic differentiations but which could also reflect immunity; and competition-colonisation trade-off, primarily conceptualised as arising from bacteriocin systems [28]. Previous work has considered the role of antigen-specific acquired immunity in maintaining strain structure [21, 26], and our insights relating to frequency-dependent epistasis are also applicable to these models. Whether similar effects are predicted to arise for other functions is, in principle, easy to address through modelling. However, the mechanisms giving rise to NFDS across the genome are not well characterised.

Thirdly, in the mechanistic models we consider, both loci are under the same form of NFDS, and it it unclear whether these effects also arise between loci under *different* forms of NFDS. Previous modelling suggests that NFDS on metabolic and antigenic loci can lead to LD between the two types of alleles [33]. In the model giving rise to these result, NFDS on antigenic and metabolic loci is assumed to arise through relatively similar mechanisms (i.e. effects on colonisation efficiency). It is unclear whether the same applies if NFDS arises through very different mechanisms for different loci. Similarly, addressing how effects combined across more than two loci under NFDS is also important.

Our results have implications for understanding the response of pathogen populations to disturbances, such as vaccinations. Associations maintained by positive frequency-dependent epistasis are contingent on the population’s evolutionary history. Such associations may therefore not be stable to large perturbations, leading to potentially restructuring of patterns of associations. We tested this in the Massachusetts data following the introduction of the pneumococcal conjugate vaccine and observed large-scale conservation of LD and few cases of LD sign reversal. More broadly, models of NFDS are used to predict which strains will increase in frequency following removal of vaccine-targetted serotypes [10, 12, 34]. While such models make considerably better predictions than naive models (e.g. *R*^2^ between observed and predicted post-vaccine strain frequencies 0.2 vs 0.02 for a NFDS model vs a model based on pre-vaccine strain frequencies), accurately predicting post-vaccine strain frequencies remains very challenging. Accounting for emerging epistatic interactions may improve the predictive power of such models.

The presence of stable strain structure requires both the presence of allelic diversity and epistatic mechanisms which maintain associations between alleles. Here, we show that NFDS, which gives rise to the former, also impacts the latter through emergent epistasis. NFDS is only neutral with respect to strain structure under very specific assumptions, and, for genes under a similar form of NFDS, there is no *a priori* reason to expect these assumptions to hold. These mechanisms that maintain genetic diversity across loci are thus also fundamentally linked to how this diversity is structured.

## Methods

### Measures of Linkage Disequilibrium

Linkage disequilibrium can be measured with the D metric defined as :

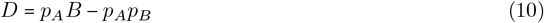

which is bounded by − 0.25 ≤ *D* ≤ 0.25. However this interval is reduced as frequencies of alleles *A* or *B* diverge from 0.5. In order to get a more interpretable metric, we use throughout most of the figures in this article the metric *D*′ which is bounded by −1 and 1 independantly of the frequencies of individual alleles (provided they are not 0 or 1):

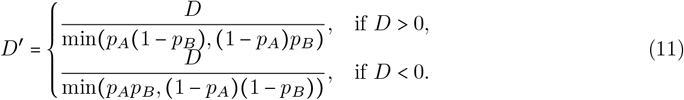

### Multi-locus NFDS

In this section we present how we implement recombination in the multi-locus NFDS model. We use a mass-action recombination where two genotypes exchange an allele at a rate dictated by their respective densities and a recombination rate *σ*. This gives the following expression for the changes in density due to recombination *φ*_*ij*_ of genotype *ij*:

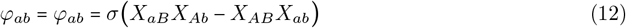

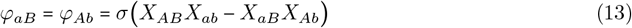

the two terms in each expression respectively showing the creation and the disappearance of the focal genotype through recombination.

### SIS with co-colonisation

We consider a SIS (uncolonised-infectious-uncolonised) for a commensal bacteria circulating in an homogeneous host population. Uncolonised hosts are characterized by a birth rate *b* and a death rate *d*. There are several strains of commensals, and we denote with *I*_*i*_ hosts colonised by strain *i*. Commensals are transmitted through a mass action mechanism with a baseline transmission rate of *β*_0_, and can be cleared with a rate *γ*. There is no virulence (or additional mortality) associated with the carriage of a commensal. In this model we allow for co-colonisation by two commensals, which can or not be the of the same strain. We denote with *I*_*i*,*j*_ such a host co-colonised by both strains *i* and *j*. Note that there is no meaning to the order of strains in the notation and so *I*_*i*,*j*_ and *I*_*j*,*i*_ can be used interchangeably. Each co-colonising bacteria is transmitted at rate *β*_0_*q* where *q* is the efficiency associated with being in co-colonisation. This means that if *q* = 1, the co-colonised host *I*_*i*,*i*_ is exactly twice as infectious as host *I*_*i*_. There is no interaction between co-colonising bacteria acting on the duration of carriage and so a co-colonised host *I*_*i*,*j*_ leaves the co-colonised compartment at rate 2*γ*: it becomes *I*_*i*_ or *I*_*j*_ at rate *γ* respectively. Two strains in co-colonisation can recombine upon secondary colonisation with probability *σ*, leading to a term Δ_*rec*_ detailed in the next section. The resulting system of ordinary differential equations is:

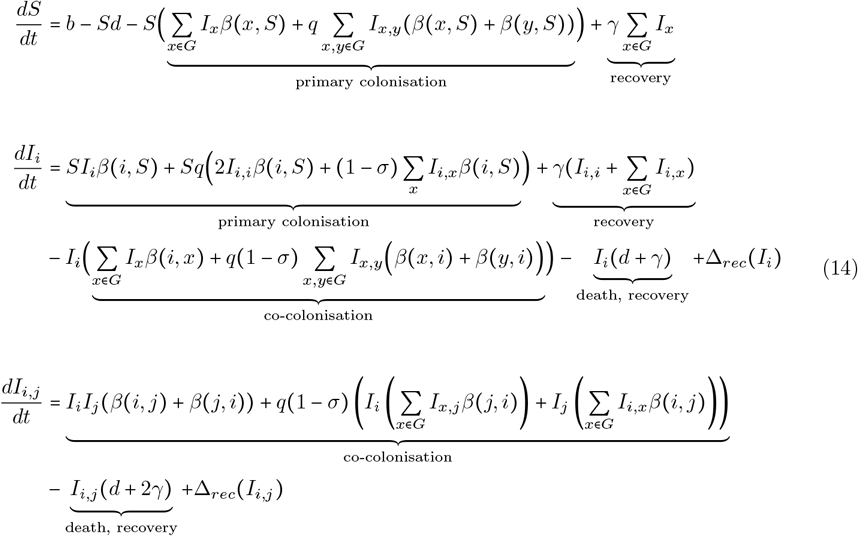

### Recombination

As we model co-colonisation, we can model recombination directly at the level of interacting strain and not use a population-wide recombination rate. We model recombination as happening upon the colonisation of a new host, from a co-colonised host. From a co-colonised host *I*_*i*,*j*_, if strain *i* recombines with strain *j* upon infection, strain *i* will receive the allele of strain *j* in a random locus. *δ*_*z*_( *i, j*) is the probability, upon recombination, that genotype *i* recombines with genotype *j* to form genotype *z*. For instance :

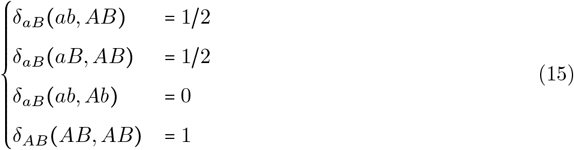

The resulting terms of growth rate dependent on recombination are then given by:

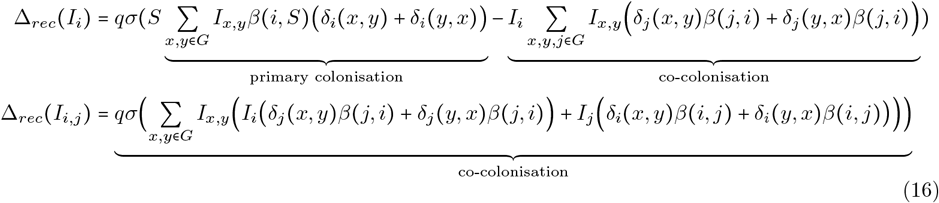

### Evolutionary dynamics of LD

Evolutionary dynamics can be monitored through the dynamics of the frequency of each allele and of linkage disequilibrium[27].

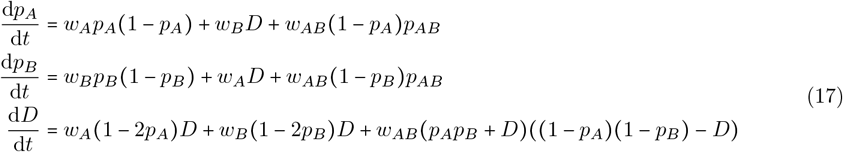

The parameter *q* is important in the expression of the epistasis *w*_*AB*_ and dictates the effective force of infection by a certain strain depending on the number of host where this strain is in single infection and the number of hosts where that strain is co-colonising. We define the effective colonising density for strain *i E*_*i*_(*t*) as:

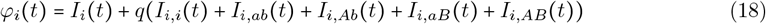

and the per-capita effective rate of new infections by strain *ab*:

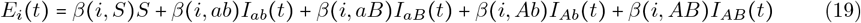

Hence *E*_*i*_ (*t*) *φ*_*i*_(*t*) is the force of infection, or rate of new colonisation by the strain *i*. With these terms we can express the growth rate of strain *i*:

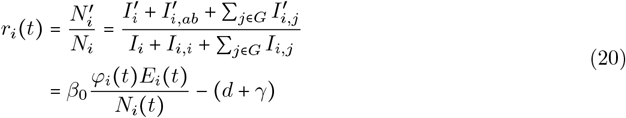

To finally get an expression for the epistasis:

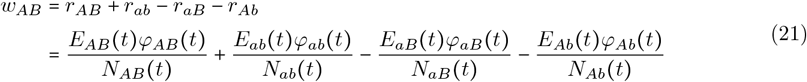

In the special case of *q* = 1, a strain in co-colonisation is as infectious as a strain in a single infected host. This lead to the equality between the effective density and the total density of strain *i φ*_*i*_(*t*) = *N*_*i*_(*t*).

### Genome data

Genomes of *S. pneumoniae* were downloaded from NCBI BioProjects (Mae La [29]:) PRJEB2357, PRJEB2393, PRJEB2395, PRJEB2479, PRJEB2480, (Southampton [30]:)PRJEB2417, (Massachusetts [31]:)PRJEB2632. The phylogenetic tree for these isolates was downloaded on https://microreact.org/project/multilocusNFDS [10]. Gene sequences were clustered into cluster of orthologous genes using the easy-cluster workflow of MMSEQS2 [35]. Ancestral reconstruction of accessory gene content was carried out with PAST-ML using the JOINT prediction method [36] and protein annotation into KEGG functional categories was done with eggnog mapper v2 [37] using the emapper workflow and the diamond database.

## Data analysis

All subsequent analysis of the genome data was carried out using R 4.3.1 [38] and the ape package [39]. Accessory genes were filtered with frequency threshold 0.1 < *f <* 0.9 in each geographical population. Next, we only kept genes that have been gained or lost on the tree more than 30 times to reduce the impact of clonal expansion and biases in the dataset.

To construct Figure 5b,c and d, we first kept only genes that are shared between two populations. Then we selected pairs of genes with an absolute LD of |*D*^′^| > 0.6 in the focal population and assessed the proportion of these gene pairs for which LD was conserved or reversed in the second population. We defined conservation as maintaining strong LD ( |*D*^′^| > 0.6) of same sign, while reversal was defined as a strong LD of opposite sign in the two populations. To assess the significance of these proportions, we built a test based on an expected distribution of these proportion using a shuffling of the second population: redistributing the gene presence/absence among the genotypes while conserving their frequencies. We performed 1000 such permutations for each test. Tests were carried out for each population pair, using each populations as the focal one. We performed one-tailed tests to test for a higher than expected number of conservation and reversal events, and two-tailed tests to test for a difference.

## Code availability

The simulation and analysis code for this paper is available at https://github.com/martingui/NFDS_epistasis.

## References

[1] Dominique A Caugant, L Oddvar Frøholm, Kjell Bøvre, Eirik Holten, Carl E Frasch, Louis F Mocca, Wendell D Zollinger and Robert K Selander. ‘Intercontinental spread of a genetically distinctive complex of clones of Neisseria meningitidis causing epidemic disease.’ In: Proceedings of the National Academy of Sciences 83.13 (1986), pp. 4927–4931.

[2] Eleanor R Watkins and Martin CJ Maiden. ‘Persistence of hyperinvasive meningococcal strain types during global spread as recorded in the PubMLST database’. In: (2012).

[3] AM Mitchell and TJ Mitchell. ‘Streptococcus pneumoniae: virulence factors and variation’. In: Clinical Microbiology and Infection 16.5 (2010), pp. 411–418.

[4] Sonja Lehtinen, François Blanquart, Nicholas J Croucher, Paul Turner, Marc Lipsitch and Christophe Fraser. ‘Evolution of antibiotic resistance is linked to any genetic mechanism affecting bacterial duration of carriage’. In: Proceedings of the National Academy of Sciences 114.5 (2017), pp. 1075–1080.

[5] Sonja Lehtinen, François Blanquart, Marc Lipsitch, Christophe Fraser and with the Maela Pneumococcal Collaboration. ‘On the evolutionary ecology of multidrug resistance in bacteria’. In: PLoS pathogens 15.5 (2019), e1007763.

[6] Michel Tibayrenc and Francisco J Ayala. ‘Reproductive clonality of pathogens: a perspective on pathogenic viruses, bacteria, fungi, and parasitic protozoa’. In: Proceedings of the National Academy of Sciences 109.48 (2012), E3305–E3313.

[7] Chrispin Chaguza, Jennifer E Cornick and Dean B Everett. ‘Mechanisms and impact of genetic recombination in the evolution of Streptococcus pneumoniae’. In: Computational and structural biotechnology journal 13 (2015), pp. 241–247.

[8] Bruce R Levin, Véronique Perrot and Nina Walker. ‘Compensatory mutations, antibiotic resistance and the population genetics of adaptive evolution in bacteria’. In: Genetics 154.3 (2000), pp. 985–997.

[9] David V McLeod and Sylvain Gandon. ‘Understanding the evolution of multiple drug resistance in structured populations’. In: Elife 10 (2021), e65645.

[10] Jukka Corander, Christophe Fraser, Michael U Gutmann, Brian Arnold, William P Hanage, Stephen D Bentley, Marc Lipsitch and Nicholas J Croucher. ‘Frequency-dependent selection in vaccine-associated pneumococcal population dynamics’. In: Nature ecology & evolution 1.12 (2017), pp. 1950–1960.

[11] Alan McNally, Teemu Kallonen, Christopher Connor, Khalil Abudahab, David M Aanensen, Carolyne Horner, Sharon J Peacock, Julian Parkhill, Nicholas J Croucher and Jukka Corander. ‘Diversification of colonization factors in a multidrug-resistant Escherichia coli lineage evolving under negative frequency-dependent selection’. In: MBio 10.2 (2019), pp. 10–1128.

[12] Taj Azarian, Pamela P Martinez, Brian J Arnold, Xueting Qiu, Lindsay R Grant, Jukka Corander, Christophe Fraser, Nicholas J Croucher, Laura L Hammitt, Raymond Reid et al. ‘Frequency-dependent selection can forecast evolution in Streptococcus pneumoniae’. In: PLoS biology 18.10 (2020), e3000878.

[13] Viggo Andreasen, Juan Lin and Simon A Levin. ‘The dynamics of cocirculating influenza strains conferring partial cross-immunity’. In: Journal of mathematical biology 35.7 (1997), pp. 825–842.

[14] Neil M Ferguson, Alison P Galvani and Robin M Bush. ‘Ecological and immunological determinants of influenza evolution’. In: Nature 422.6930 (2003), pp. 428–433.

[15] Sarah Cobey. ‘Pathogen evolution and the immunological niche’. In: Annals of the New York Academy of Sciences 1320.1 (2014), pp. 1–15.

[16] Jun Feng, Yili Qian, Zhichao Zhou, Sarah Ertmer, Eugenio I Vivas, Freeman Lan, Joshua J Hamilton, Federico E Rey, Karthik Anantharaman and Ophelia S Venturelli. ‘Polysaccharide utilization loci in Bacteroides determine population fitness and community-level interactions’. In: Cell host & microbe 30.2 (2022), pp. 200–215.

[17] George Livingston, Miguel Matias, Vincent Calcagno, Claire Barbera, Marine Combe, Mathew A Leibold and Nicolas Mouquet. ‘Competition–colonization dynamics in experimental bacterial metacommunities’. In: Nature communications 3.1 (2012), p. 1234.

[18] Dustin Brisson. ‘Negative frequency-dependent selection is frequently confounding’. In: Frontiers in Ecology and Evolution 6 (2018), p. 10.

[19] Gabrielle L Harrow, John A Lees, William P Hanage, Marc Lipsitch, Jukka Corander, Caroline Colijn and Nicholas J Croucher. ‘Negative frequency-dependent selection and asymmetrical transformation stabilise multi-strain bacterial population structures’. In: The ISME Journal 15.5 (2021), pp. 1523–1538.

[20] Sarah Cobey and Marc Lipsitch. ‘Niche and neutral effects of acquired immunity permit coexistence of pneumococcal serotypes’. In: Science 335.6074 (2012), pp. 1376–1380.

[21] Sunetra Gupta, Martin CJ Maiden, Ian M Feavers, Sean Nee, Robert M May and Roy M Anderson. ‘The maintenance of strain structure in populations of recombining infectious agents’. In: Nature medicine 2.4 (1996), pp. 437–442.

[22] M Gabriela M Gomes, Graham F Medley and D James Nokes. ‘On the determinants of population structure in antigenically diverse pathogens’. In: Proceedings of the Royal Society of London. Series B: Biological Sciences 269.1488 (2002), pp. 227–233.

[23] Mario Recker and Sunetra Gupta. ‘A model for pathogen population structure with cross-protection depending on the extent of overlap in antigenic variant repertoires’. In: Journal of theoretical biology 232.3 (2005), pp. 363–373.

[24] Caroline O Buckee, Keith A Jolley, Mario Recker, Bridget Penman, Paula Kriz, Sunetra Gupta and Martin CJ Maiden. ‘Role of selection in the emergence of lineages and the evolution of virulence in Neisseria meningitidis’. In: Proceedings of the National Academy of Sciences 105.39 (2008), pp. 15082–15087.

[25] Shai Pilosof, Qixin He, Kathryn E Tiedje, Shazia Ruybal-Pesántez, Karen P Day and Mercedes Pascual. ‘Competition for hosts modulates vast antigenic diversity to generate persistent strain structure in Plasmodium falciparum’. In: PLoS biology 17.6 (2019), e3000336.

[26] David V McLeod, Claudia Bank and Sylvain Gandon. ‘A multilocus perspective on the evolutionary dynamics of multistrain pathogens’. In: Proceedings of the National Academy of Sciences 121.42 (2024), e2401578121.

[27] Troy Day and Sylvain Gandon. ‘The evolutionary epidemiology of multilocus drug resistance’. In: Evolution 66.5 (2012), pp. 1582–1597.

[28] Sonja Lehtinen, Nicholas J Croucher, François Blanquart and Christophe Fraser. ‘Epidemiological dynamics of bacteriocin competition and antibiotic resistance’. In: Proceedings of the Royal Society B 289.1984 (2022), p. 20221197.

[29] Claire Chewapreecha, Simon R Harris, Nicholas J Croucher, Claudia Turner, Pekka Marttinen, Lu Cheng, Alberto Pessia, David M Aanensen, Alison E Mather, Andrew J Page et al. ‘Dense genomic sampling identifies highways of pneumococcal recombination’. In: Nature genetics 46.3 (2014), pp. 305–309.

[30] Rebecca A Gladstone, Vanessa Devine, Jessica Jones, David Cleary, Johanna M Jefferies, Stephen D Bentley, Saul N Faust and Stuart C Clarke. ‘Pre-vaccine serotype composition within a lineage signposts its serotype replacement–a carriage study over 7 years following pneumococcal conjugate vaccine use in the UK’. In: Microbial Genomics 3.6 (2017), e000119.

[31] Nicholas J Croucher, Jonathan A Finkelstein, Stephen I Pelton, Patrick K Mitchell, Grace M Lee, Julian Parkhill, Stephen D Bentley, William P Hanage and Marc Lipsitch. ‘Population genomics of post-vaccine changes in pneumococcal epidemiology’. In: Nature genetics 45.6 (2013), pp. 656– 663.

[32] Aswin Krishna, Gerry Tonkin-Hill, Thibaut Morel-Journel, Stephen Bentley, Paul Turner, François Blanquart and Sonja Lehtinen. ‘Quantifying the effects of antibiotic resistance and within-host competition on strain fitness in Streptococcus pneumoniae’. In: bioRxiv (2025), pp. 2025–03.

[33] Eleanor R Watkins, Bridget S Penman, José Lourenço, Caroline O Buckee, Martin CJ Maiden and Sunetra Gupta. ‘Vaccination drives changes in metabolic and virulence profiles of Streptococcus pneumoniae’. In: PLoS pathogens 11.7 (2015), e1005034.

[34] Caroline Colijn, Jukka Corander and Nicholas J Croucher. ‘Designing ecologically optimized pneumococcal vaccines using population genomics’. In: Nature Microbiology 5.3 (2020), pp. 473– 485.

[35] Martin Steinegger and Johannes Söding. ‘MMseqs2 enables sensitive protein sequence searching for the analysis of massive data sets’. In: Nature biotechnology 35.11 (2017), pp. 1026–1028.

[36] Sohta A Ishikawa, Anna Zhukova, Wataru Iwasaki and Olivier Gascuel. ‘A fast likelihood method to reconstruct and visualize ancestral scenarios’. In: Molecular biology and evolution 36.9 (2019), pp. 2069–2085.

[37] Carlos P Cantalapiedra, Ana Hernández-Plaza, Ivica Letunic, Peer Bork and Jaime Huerta-Cepas. ‘eggNOG-mapper v2: functional annotation, orthology assignments, and domain prediction at the metagenomic scale’. In: Molecular biology and evolution 38.12 (2021), pp. 5825–5829.

[38] R Core Team. R: A Language and Environment for Statistical Computing. R Foundation for Statistical Computing. Vienna, Austria, 2023.

[39] Emmanuel Paradis and Klaus Schliep. ‘ape 5.0: an environment for modern phylogenetics and evolutionary analyses in R’. In: Bioinformatics 35 (2019), pp. 526–528.

