## Supplementary Information for "Emergent epistasis mediates the role of negative frequency-dependent selection in bacterial strain structure"

### Emergent epistasis mediates the role of NFDS in bacterial strain structure: Supplementary Information

#### Supplementary Note 1: One metabolic locus with two alleles

In this note we describe explicitly the dynamical equations for all compartments of the model in the case of one metabolic locus with two alleles. In this case efficiency of co-colonisation is only $k_0$ if the allele of the co-colonising and resident strain is the same or $k_1$ if the two alleles are different.

$$
\begin{aligned}
\frac{dS}{dt} &= b - Sd - S\beta_0\Big((I_a + I_A) + 2q(I_{a,a} + I_{A,a} + I_{A,A})\Big) + \gamma(I_a + I_A) \\
\frac{dI_a}{dt} &= S\beta_0 I_a + S\beta_0 q(2I_{a,a} + I_{A,a}) \\
&+ \gamma(2I_{a,a} + I_{a,A}) - I_a(d + \gamma) \\
&- \beta_0 I_a(k_1 I_a + k_0 I_A) - \beta_0 I_a q\,(2k_1 I_{a,a} + (k_0 + k_1)I_{a,A} + 2k_0 I_{A,A}) \\
\frac{dI_A}{dt} &= S\beta_0 I_A + S\beta_0 q(2I_{A,A} + I_{a,A}) \\
&+ \gamma(2I_{A,A} + I_{a,A}) - I_A(d + \gamma) \\
&- \beta_0 I_A(k_1 I_A + k_0 I_a) - \beta_0 I_A q\,(2k_1 I_{A,A} + (k_0 + k_1)I_{a,A} + 2k_0 I_{a,a}) \\
\frac{dI_{a,a}}{dt} &= 2\beta_0 I_a^2 - I_{a,a}(d + 2\gamma) \\
&+ \beta_0 q I_a k_1(2I_{a,a} + I_{a,A}) \\
\frac{dI_{A,A}}{dt} &= 2\beta_0 I_A^2 - I_{A,A}(d + 2\gamma) \\
&+ \beta_0 q I_A k_1(2I_{A,A} + I_{a,A}) \\
\frac{dI_{a,A}}{dt} &= 2\beta_0 I_a I_A - I_{a,A}(d + 2\gamma) \\
&+ \beta_0 q\big(I_a k_0(2I_{A,A} + I_{a,A}) + I_A k_0(2I_{a,a} + I_{a,A})\big)
\end{aligned}
\tag{1}
$$

Note that recombination terms vanish in this system with only one locus when recombination is symmetric. This would not be the case if recombination was skewed. For instance if allele a and A were respectively the absence and the presence of a gene, one could consider that the gene is preferentially lost rather than acquired, in this case the asymmetric recombination term would remain.

To highlight the negative frequency-dependent selection (NFDS) in this system we introduce

$$
\begin{aligned}
p_A &= \frac{I_A + I_{a,A} + 2I_{A,A}}{I_a + I_A + 2I_{a,a} + 2I_{a,A} + 2I_{A,A}} \\
&= \frac{N_A}{N_{tot}}
\end{aligned}
\tag{2}
$$

the frequency of allele A in the bacterial population, with $N_A$ the density of bacteria of strain A and $N_{tot}$ the total density of bacteria. The dynamics of this frequency can be derived as a function of the selection coefficient for allele A which is defined by

$$w_A(t) = r_A(t) - r_a(t) \tag{3}$$

where $r_A(t)$ and $r_a(t)$ are respectively the growth rate of bacteria with allele A and a in the population such that

$$\begin{aligned} r_A(t) &= \frac{N'_A}{N_A} = \frac{I'_A + 2I'_{A,A} + I'_{a,A}}{I_A + I_{a,A} + 2I_{A,A}} \\ r_a(t) &= \frac{N'_a}{N_a} = \frac{I'_a + 2I'_{a,a} + I'_{a,A}}{I_a + I_{a,A} + 2I_{a,a}} \end{aligned} \tag{4}$$

We get:

$$\frac{\mathrm{d}p_A}{\mathrm{d}t} = w_A p_A (1 - p_A) \tag{5}$$

First when $q = 1$, meaning that a bacteria in a co-colonised host is as infectious as a bacteria in a single-infected host (and thus $I_{A,A}$ is twice as infectious as $I_A$), we get

$$w_A(t) = \beta_0 (k_0 - k_1)(I_a - I_A) \tag{6}$$

We see with this equation when $k_0 > k_1$, which is true in a context of metabolic niche differentiation, that the selection coefficient for allele A is positive when $I_A < I_a$. Hence the less frequent strain will have a higher growth rate than the more frequent strain, and the system is only at stable equilibrium when $I_A = I_a$.

We also derive this selection coefficient for the special value of $q = 1/2$, meaning that a bacteria in a co-colonised host is half as infectious as a bacteria in a single infected host (and so $I_{A,A}$ is equally as infectious as $I_A$). In this case we get

$$\begin{aligned} w_A(t) &= \frac{1}{2}\beta_0 \Big( (k_0 - k_1)(I_a - I_A) + \frac{I_A(S + k_0 I_a + k_1 I_A)}{I_A + I_{a,A} + 2I_{A,A}} - \frac{I_a(S + k_0 I_A + k_1 I_a)}{I_a + I_{a,A} + 2I_{a,a}} \Big) \\ &= \frac{1}{2}\beta_0 \Big( (k_0 - k_1)(I_a - I_A) + \frac{I_A(S + k_0 I_a + k_1 I_A)}{N_A} - \frac{I_a(S + k_0 I_A + k_1 I_a)}{N_a} \Big) \end{aligned} \tag{7}$$

Again we find a term like in the previous equation with the difference between the density of both single-infected types which lead to NFDS. The next terms can be seen as the rate of new transmission produced by the single infected of each type, divided by the total density of that strain in the population.

To get a better understanding of these last two terms, one can look at the expression of the selection coefficient when $q = 0$, meaning a strain in co-colonisation is able to transmit:

$$w_A(t) = \beta_0\Big(\frac{I_A(S + k_0 I_a + k_1 I_A)}{N_A} - \frac{I_a(S + k_0 I_A + k_1 I_a)}{N_a}\Big) \tag{8}$$

In that case we find a very different behaviour (see Figure S2). For $q = 0$, only the bacteria in a single infected host can transmit, which means that the rate at which it co-colonises has not direct effect on the selection coefficient ie. on fitness. However, if only single infected hosts can transmit and only the primary colonisation affect the fitness directly, the driver of the fitness difference between the strains becomes the ability for resident strains to avoid being co-colonised. Thus if the likelihood of being co-colonised increases, fitness decreases. In our metabolic niche framework, a strain is more easily co-colonised by a strain with a different allele. This create a positive frequency-dependent selection on allele frequency: if an allele is in higher frequency, the bacteria carrying this allele will be more likely to evade being co-colonised, and their frequency will increase. Thus going back to the expression of $w_A$ for the more realistic value of $q = 1/2$, we see a combination of the two contrasting effects that are clearly highlighted for the values $q = 0$ or $q = 1$. Figure S2 highlights how we find a transition between NFDS and PFDS due to these effects for a value close to $q = 0.1$.

### Supplementary Note 2: Competition-colonisation with one bi-allelic locus

We now describe explicitly the system of ODEs for the competition-colonisation trade-off model for one bi-allelic locus. In this model, $m$ is the benefit of the coloniser allele $a$ on the primary colonisation. $k_{-1} < k_0 < k_{+1}$ represent the efficiency of co-colonisation depending on the whether the coloniser has respectively one more, the same number, or one less competitive allele A than the resident strain.

$$\begin{aligned}
\frac{dS}{dt} &= b - dS + \gamma(I_a + I_A) - S\beta_0\Big((1+m)I_a + I_A + q(2(1+m)I_{a,a} + (2+m)I_{a,A} + 2I_{A,A})\Big) \\
\frac{dI_a}{dt} &= \beta_0(1+m)SI_a + \beta_0 Sq\left(2(1+m)I_{a,a} + (1+m)I_{a,A}\right) \\
&\quad + \gamma\left(2I_{a,a} + I_{a,A}\right) - I_a(d+\gamma) \\
&\quad - \beta_0 I_a\left(k_0 I_a + k_{+1} I_A\right) - q\beta_0 I_a\left(2k_0 I_{a,a} + (k_0 + k_{+1})I_{a,A} + 2k_{+1} I_{A,A}\right) \\
\frac{dI_A}{dt} &= \beta_0 SI_A + S\beta_0 q\left(I_{a,A} + 2I_{A,A}\right) \\
&\quad + \gamma\left(I_{a,A} + 2I_{A,A}\right) - I_A(d+\gamma) \\
&\quad - \beta_0 I_A\left(k_{-1} I_a + k_0 I_A\right) - q\beta_0 I_A\left(2k_{-1} I_{a,a} + (k_{-1} + k_0)I_{a,A} + 2k_0 I_{A,A}\right) \\
\frac{dI_{a,a}}{dt} &= \beta_0 k_0 I_a^2 + q\beta_0 k_0 I_a\left(2I_{a,a} + I_{a,A}\right) - (d+2\gamma)I_{a,a} \\
\frac{dI_{A,A}}{dt} &= \beta_0 k_0 I_A^2 + q\beta_0 k_0 I_A\left(I_{a,A} + 2I_{A,A}\right) - (d+2\gamma)I_{A,A} \\
\frac{dI_{a,A}}{dt} &= \beta_0(k_{-1} + k_{+1})I_a I_A - (d+2\gamma)I_{a,A} \\
&\quad + q\beta_0\left[I_A\left(2k_{-1} I_{a,a} + k_{-1} I_{a,A}\right) + I_a\left(k_{+1} I_{a,A} + 2k_{+1} I_{A,A}\right)\right]
\end{aligned} \tag{9}$$

With the same reasoning as the previous section, we get for the case $q = 1$ the expression for the selection coefficient of allele A as:

$$w_A(t) = \beta_0\left(\left(k_{+1} - k_0\right) I_0(t) + \left(k_0 - k_{-1}\right) I_1(t) - mS(t)\right) \tag{10}$$

Therefore, selection favours the coloniser allele a when the number of un-colonised hosts $S$ is high compared to the number of single colonised hosts $I_a$ and

$I_A$. However, when the coloniser allele increases in frequency, there is a stronger pressure and thus a depletion of $S$ hosts compared to single colonised hosts. In turn, this favours the selection for the competitive allele, thus leading to balancing selection. However, the equilibrium frequency of each allele is not necessarily 0.5, and diversity is not necessarily maintained for all sets of parameters. We highlight this in Figure S6a and S6b, showing how the equilibrium frequency of the competitive allele $A$ increases with $k_{+1}$ and decreases with $m$ or $k_{-1}$.

Similarly to the previous model, we also explore the case where $q = 0.5$, meaning that the efficiency of colonising from a co-colonised host is reduced by half compared to a single colonised host. The effect on the selection coefficient is

$$\begin{aligned} w_A(t) = \frac{\beta_0}{2}\Big( & (k_{+1} - k_0)\, I_0(t) + (k_0 - k_{-1})\, I_1(t) - mS(t) \\ & + \frac{I_A(S + k_{+1}I_a + k_0 I_A)}{N_A} - \frac{I_a(S + k_{-1}I_A + k_0 I_a)}{N_a}\Big) \end{aligned} \tag{11}$$

Balancing selection therefore emerges as the density of uncolonised hosts reduces the selection coefficient for the competitive genotype $A$, which depletes the $S$ population less than the coloniser $a$. The impact on the frequency of the competitive allele is presented in Figure S6c and S6d.

#### Supplementary Note 3: Exploring the outcomes of the competition-colonisation model

To explore the outcomes of the competition-colonisation model (Figure S7 and S8), we use two meta-parameters to explore different geometries for the competition efficiencies $k_\Delta$. We use a concavity parameter $p_c$ and a step parameter $p_s$, which we implement as:

$$\begin{aligned} k_{-1} &= logistic(logit(0.25) + p_c + 0.5p_s) \\ k_0 &= logistic(logit(0.5) + p_c - p_s) \\ k_{+1} &= logistic(logit(0.75) + p_c + 0.5p_s) \end{aligned} \tag{12}$$

using the logistic and logit functions defined as :

$$logistic(x) = \frac{1}{1+e^{-x}}$$
$$logit(x) = Log(\frac{x}{1-x}) \tag{13}$$

The effects of these parameters on the geometries of the $k$ parameters and on equilibrium LD are shown both in Figure S7 and S8 respectively in the absence and presence of recombination.

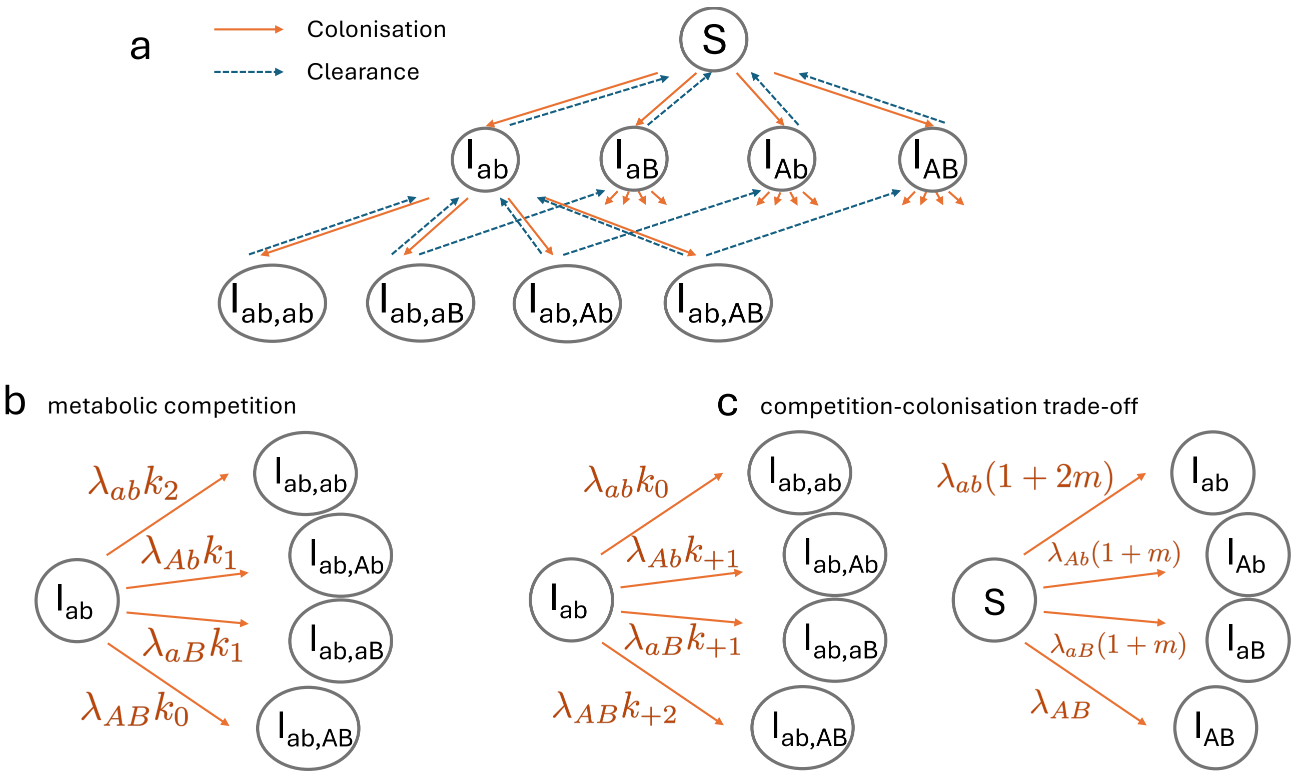


**Figure S1: Co-colonisation framework with two bi-allelic loci.** (a) We represent a schematic of the co-colonisation framework we use throughout this work. Hosts can be uncolonised ($S$), single colonised ($I_x$) or co-colonised ($I_{x,y}$). Hosts can be co-colonised by the same strain twice. In the metabolic niche model (b), co-colonisation rates are reduced when the resident and incoming strains share the same allele, modeled by parameters $k_2 < k_1 < k_0$. We show the rates of colonisation for a focal host already colonised by strain $ab$. In the competition-colonisation model (c), lower case alleles are "colonising", leading to an increased rate of primary colonisation $m$, while the competitive upper case alleles leads to higher rates of co-colonisation on average. Co-colonisation rates are dependent on the difference in number of competitive alleles between the incoming and the resident strain $\Delta$ through the parameters $k_\Delta$. Again, we show the rates of colonisation for a focal host already colonised by strain $ab$. $\lambda_{ab} = \beta_0(I_{ab} + q(I_{ab,ab} + \sum_{j\in G} I_{ab,j})$ represents the partial force of infection of genotype $ab$, taking into account that strains in co-colonisation have reduced transmission efficiency through parameter $q$. Solid orange lines represent colonisation and dashed blue lines represent clearance.

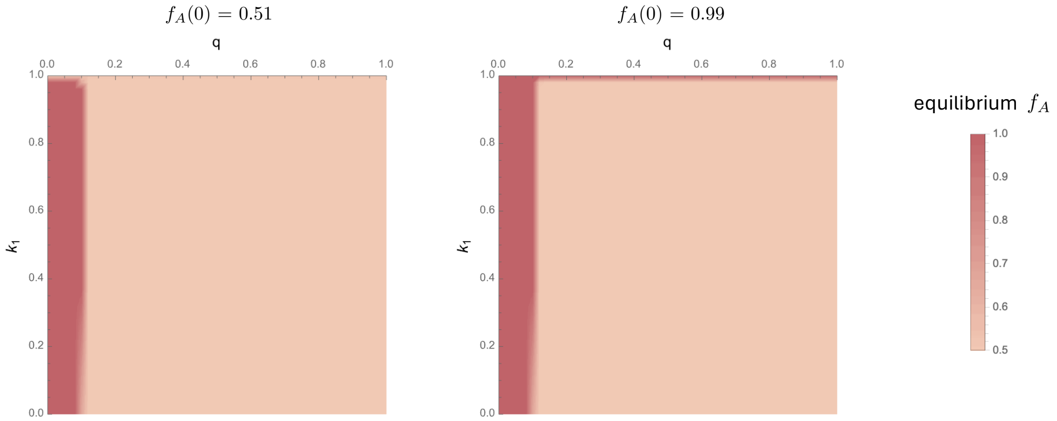


**Figure S2: Equilibrium frequency in the one locus metabolic niche model** as a function of competition parameter $k_1$ and efficiency of transmitting from a co-colonised host $q$, for different starting frequencies $p_A$. Note that the behaviour for a starting frequency $p_A < 0.5$ is symmetric, due to the symmetry of alleles a and A. Parameter values used: $\beta 0 = 2$, $b = 4$, $\gamma = 2$, $d = 1$, $k_0 = 1$, $q = 0.5$.

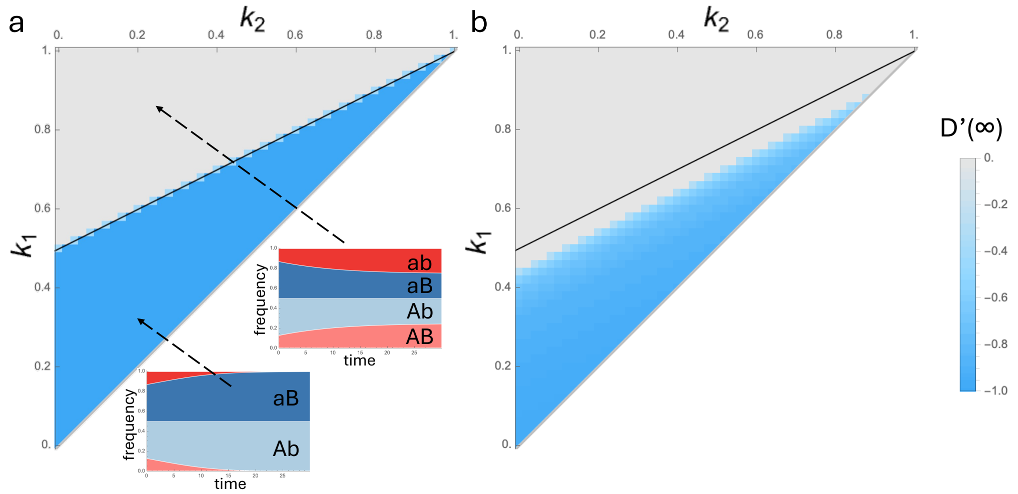


**Figure S3: Equilibrium LD is null of negative when starting with an initially negative LD.** This figure reproduces the results of Figure 2 but with a negative starting value of linkage disequilibrium D(0). Mller plots in panel (a) show the dynamics of the cumulative frequencies of all genotypes through time, depending on the competition parameters, starting for an initial LD of $D = -0.1$. Parameter values used: $\beta 0 = 2$, $b = 4$, $\gamma = 2$, $d = 1$, $k_0 = 1$, $q = 0.5$.

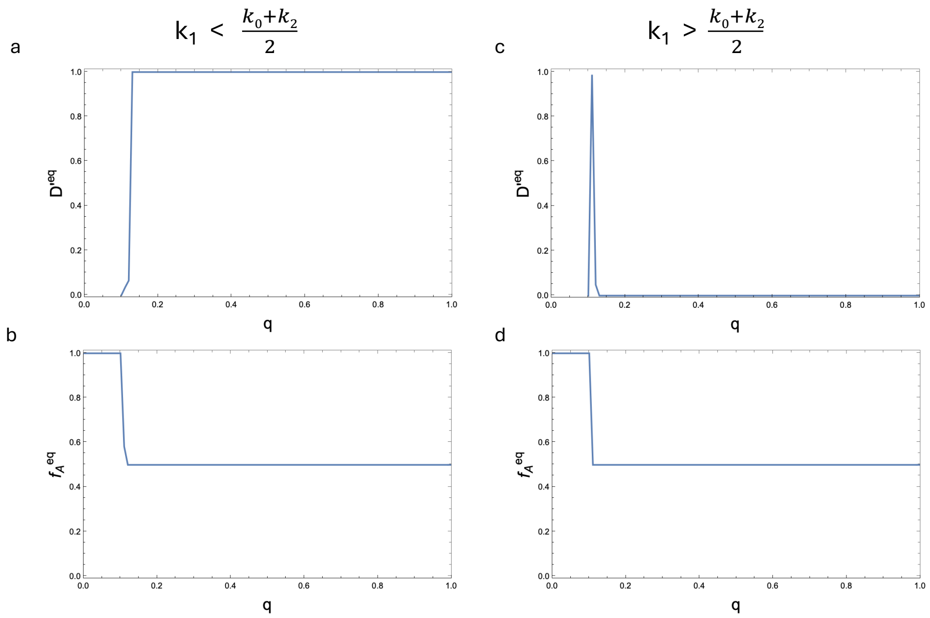


**Figure S4: Equilibrium LD (a,c) and allele frequency (b,d) in the metabolic niche model as a function of $q$.** Values of equilibrium $D'$ are not shown when $f_A = 1$ because it is not defined. Initial LD is $D' = 0.2$. We chose initial values of frequencies $f_A(0) = f_B(0) = 0.501$ to avoid numerical instabilities. For panels (a,b): $k_2 = 0.2, k_1 = 0.5, k_0 = 1$, or panels (c,d): $k_2 = 0.2, k_1 = 0.7, k_0 = 1$. Other parameter values used: $\beta 0 = 2$, $b = 4$, $\gamma = 2$, $d = 1$.

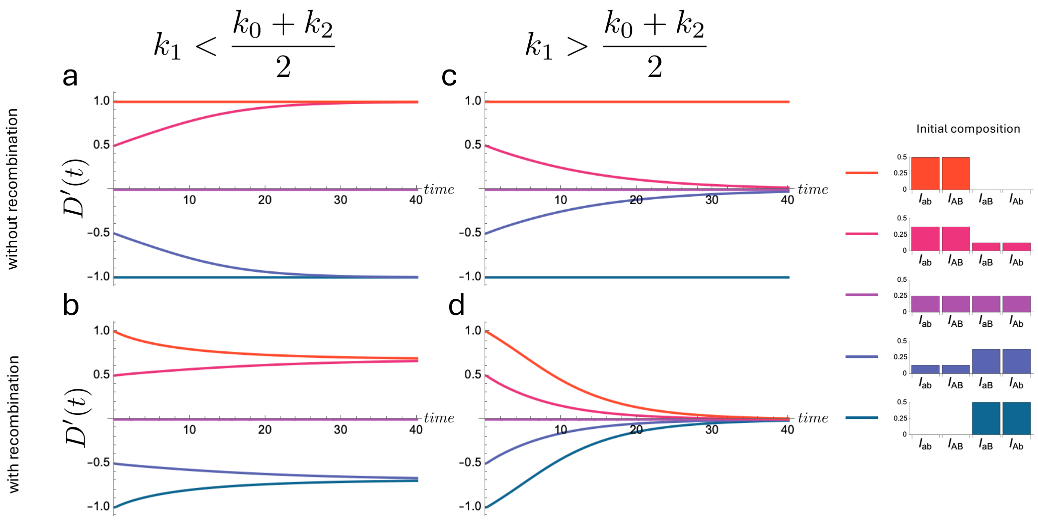


**Figure S5: Linkage disequilibrium trajectories in the two different competitive regimes of the metabolic niche model: NFDS or PFDS.** The values of LD are shown through time for 5 different initial population compositions, shown on the right, characterised by values of initial LD of $D' \in \{-1, -0.5, 0, 0.5, 1\}$. For panels (a,b): $k_2 = 0.2, k_1 = 0.5, k_0 = 1$, for panels (c,d): $k_2 = 0.2, k_1 = 0.7, k_0 = 1$. Other parameter values used: $\beta 0 = 2$, $b = 4$, $\gamma = 2$, $d = 1$, $q = 0.5$.

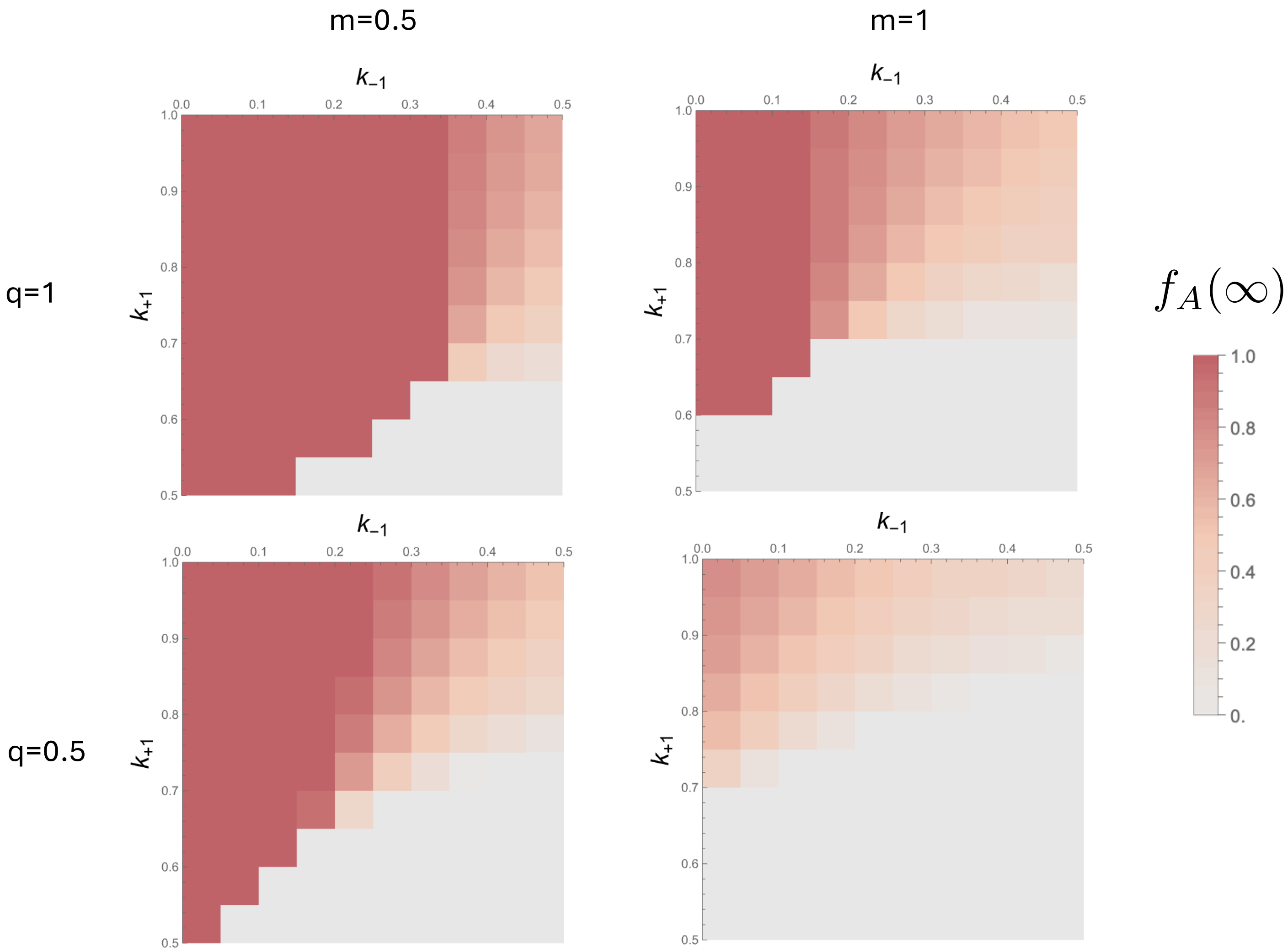


**Figure S6: Equilibrium frequency of the competitive allele $A$ in the competition-colonisation model** according to the competitive efficiencies $k_{-1}$ and $k_{+1}$, with different values of $q$ the efficiency of colonisation from a co-colonised host and the benefit of the colonising allele $m$. Parameter values used: $\beta 0 = 2$, $b = 4$, $\gamma = 2$, $d = 1\, \sigma = 0$, $k_0 = 0.5$.

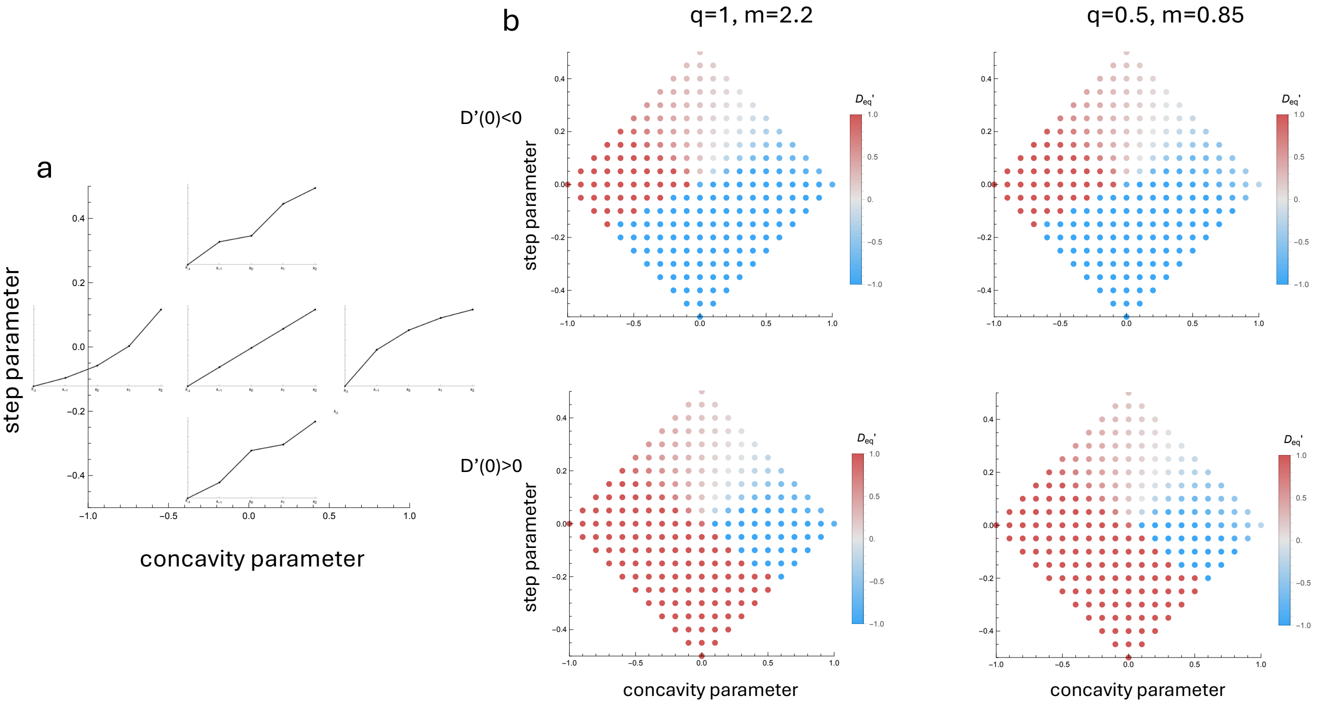


**Figure S7: Strain structuring effects of a competition-colonisation trade-off model.** The outcomes of the models are explored according to two meta-parameters that control the geometry of the competition efficiencies (a), a concavity parameter and a step parameter (see Methods, eq. (12)). (b) Equilibrium $D'$ is shown as a function of the step and concavity parameters starting from a negative (first row) or a positive (second row) value of linkage disequilibrium $D'(0)$. The two columns show the results for two values of efficiency from a co-colonised host $q$ and colonising benefit $m$. The value of $m$ was chosen so that in the additive case, the frequency of each allele was close to 0.5. Recombination is absent. For the case with recombination, see Figure S8. Parameter values used: $\beta 0 = 2$, $b = 4$, $\gamma = 2$, $d = 1\, \sigma = 0$.

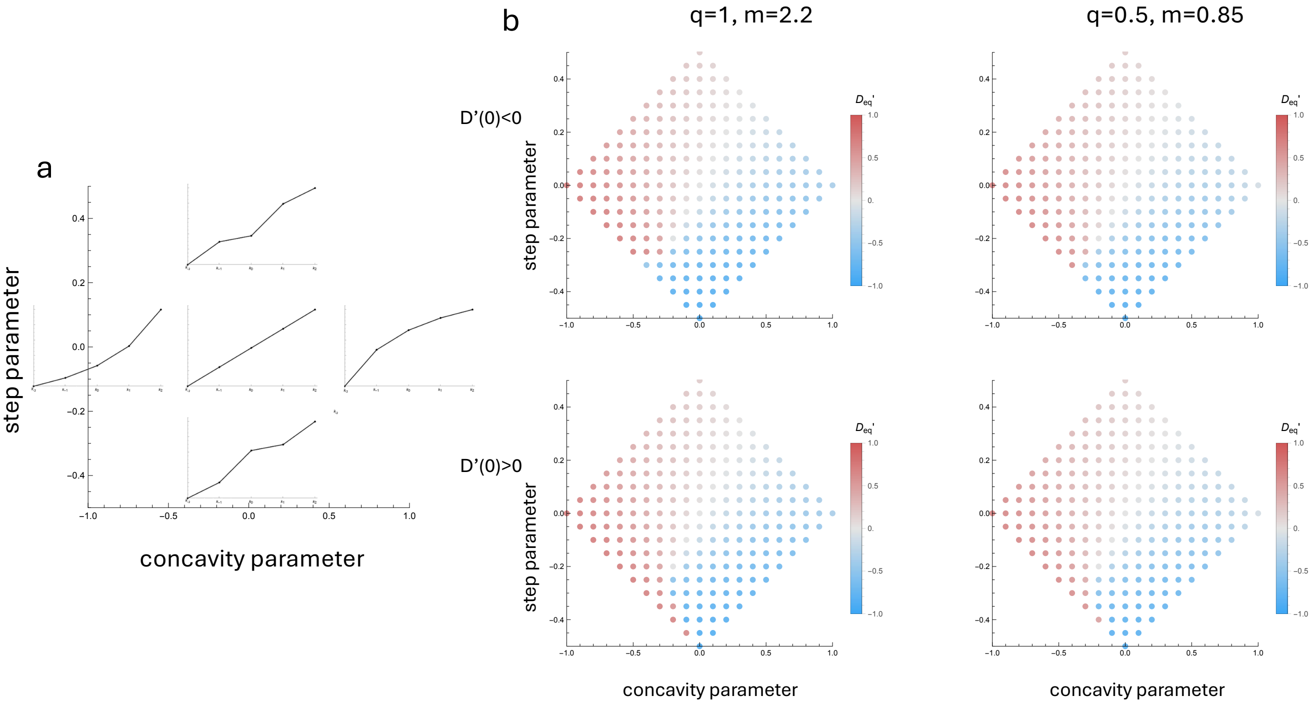


**Figure S8: Strain structuring effects of a competition-colonisation trade-off model in the presence of recombination.** The outcomes of the models are explored according to two meta-parameters that control the geometry of the competition efficiencies (a), a concavity parameter and a step parameter (see Methods, eq. (12)). (b) Equilibrium $D'$ is shown as a function of the step and concavity parameters starting from a negative (first row) or a positive (second row) value of linkage disequilibrium $D'(0)$. The two columns show the results for two values of efficiency from a co-colonised host $q$ and colonising benefit $m$. The value of $m$ was chosen so that in the additive case, the frequency of each allele was close to 0.5. Parameter values used: $\beta 0 = 2$, $b = 4$, $\gamma = 2$, $d = 1$, $\sigma = 0.05$.

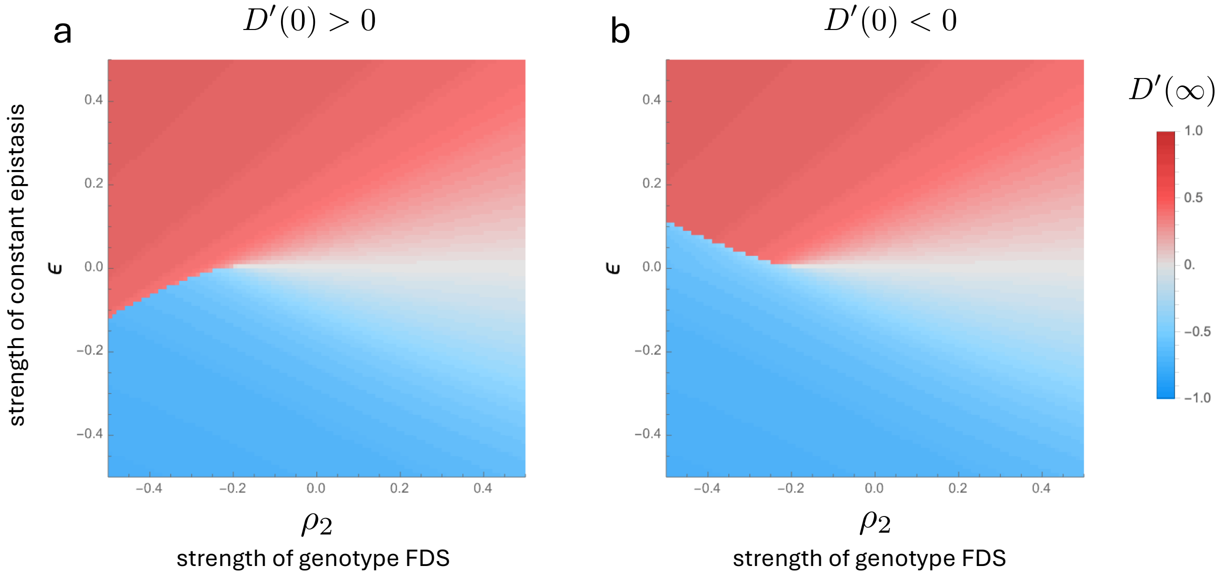


**Figure S9: The simple multi-locus NFDS model with epistasis and recombination.** Equilibrium $D'$ is shown as a function of the constant epistasis parameter $\epsilon$ and genotype frequency-dependent epistasis parameter $\rho_2$ starting from a positive (a) or a negative (b) linkage disequilibrium $D'(0)$, in the presence of recombination. Parameter values used: $r = 1,\ \rho = 1,\ \kappa = 1, \sigma = 0.05,\ p^*_A = p^*_B = 0.5,\ \ p^*_{ij} = 0.25.$

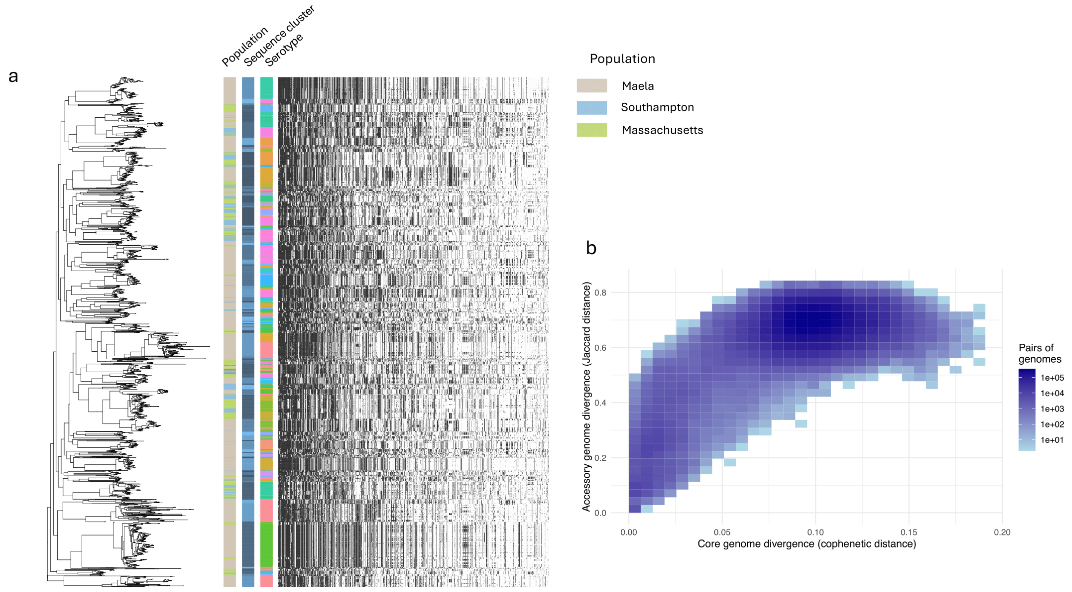


**Figure S10: The strain structure of *Streptococcus pneumoniae*.** (a) Distribution of sequence clusters, serotypes and accessory genes on the phylogenetic tree of *Streptococcus pneumoniae*. Different colours indicate different sequence clusters and serotypes. Accessory genes are sorted left to right from most frequent to less frequent. Only genes with frequency between 0.1 and 0.9, and with more than 30 gains and losses throughout the tree are shown. (b) Gene content relatedness as a function of phylogenetic distance for all pairs of genomes.

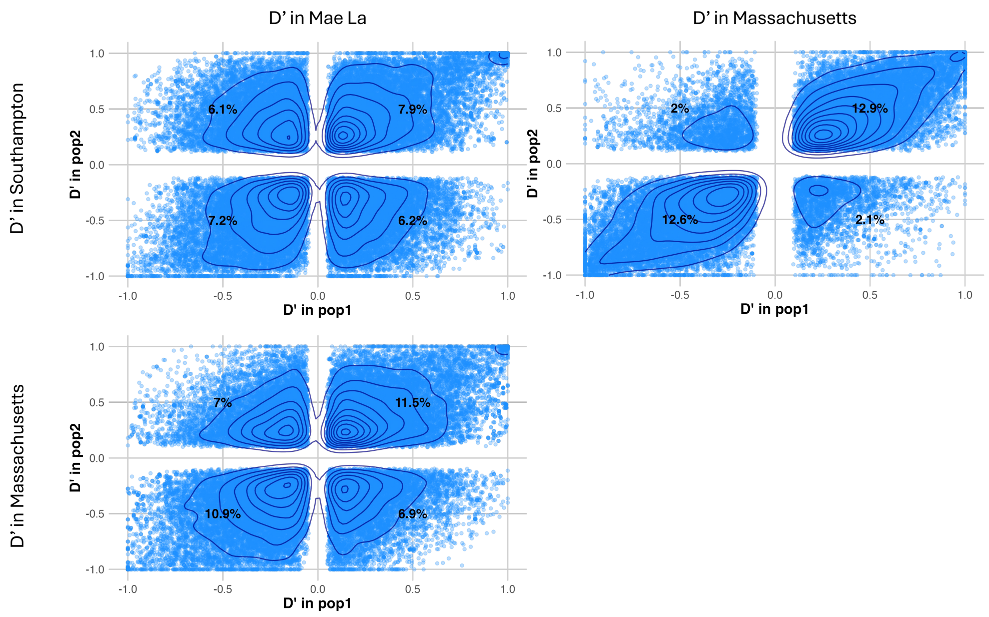


**Figure S11: Comparison of $D'$ for all pairs of genes across populations.** Only pairs with a significant linkage, as assessed with a chi-square, B-H corrected p-value<0.05 are shown. The percentage values reflect the percentage of pairs in that quadrant.

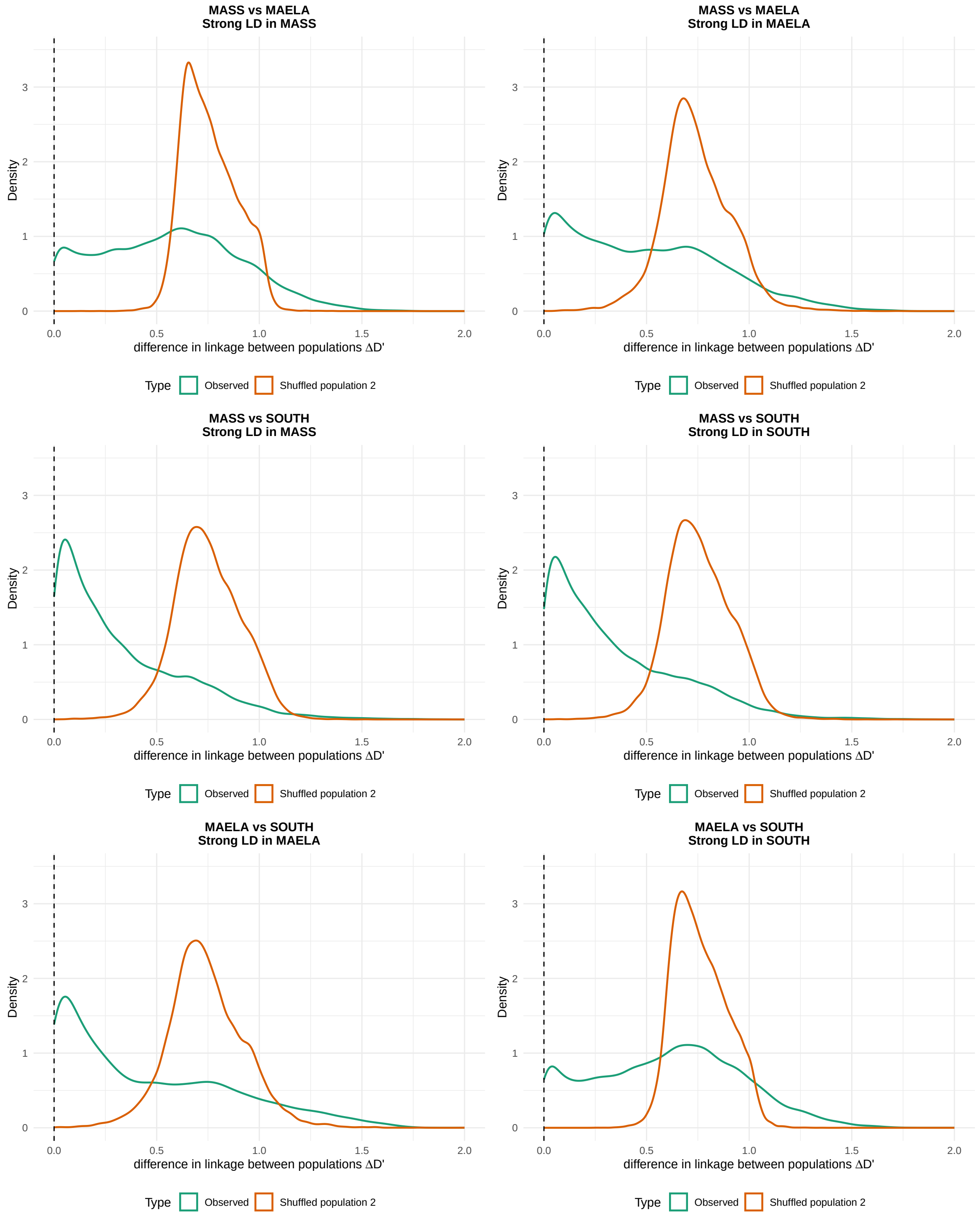


**Figure S12: Extension of main Figure 4 including all pairs of population, and all focal populations.** For instance the first row compares the Massachusetts and MaeLa datasets. The left plot focuses on genes strongly linked in the Massachusetts dataset, with or without permutating the MaeLa dataset. The plot on the right focuses on genes strongly linked in the MaeLa dataset, permutating or not the Massachusetts dataset.

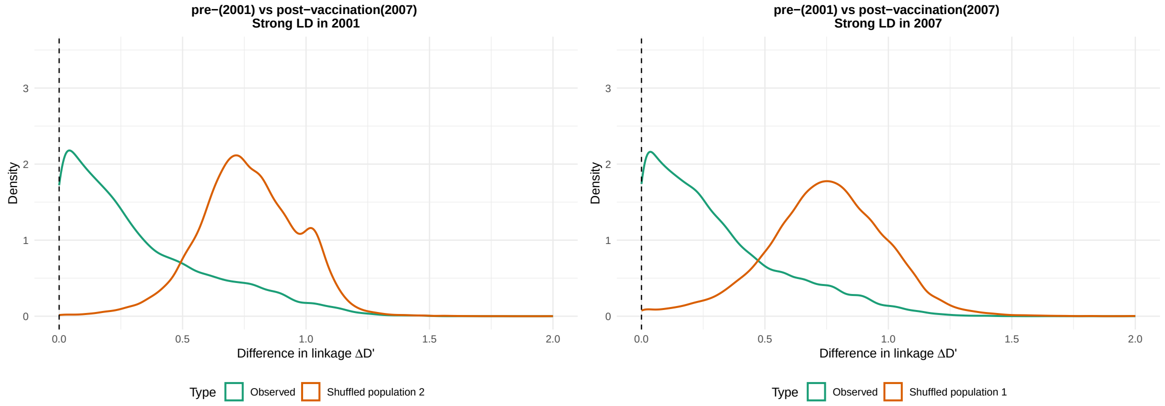


**Figure S13: Patterns of LD change across time in Massachusetts between 2001 (before vaccination) and 2007 (after vaccination) for each pair of gene with strong LD in the 2001 dataset.** We study the distributions of absolute difference of LD across populations. We show for gene pairs highly linked in the 2001 dataset ($|D'| > 0.6$) the distribution of change in linkage compared to the 2007 dataset. The green line shows the distribution of observed changes, and the orange line shows the distribution when the gene presence/absence data is permuted in the 2007 population.
